# *Drosophila* Metaxin-2 controls beta-barrel protein biogenesis and muscle growth in a stage-dependent fashion

**DOI:** 10.1101/2025.05.22.655489

**Authors:** Xinyi Shou, Xiaoyu Fan, Ling Li, Yina Ruan, Wei Li, Weina Shang, Jianhua Mao, Xiaojun Xie

## Abstract

Metaxin-2 (Mtx2) is an evolutionarily conserved mitochondrial outer membrane protein. Mutations in human *Mtx2* cause mandibuloacral dysplasia (MADaM), a progeroid disorder. However, the pathologic mechanisms of *Mtx2* loss-of-function remain largely unknown. Using *Drosophila*, we show *Mtx2* null mutants exhibit pupal lethality, rescued by *Drosophila* or human Mtx2, underscoring functional conservation across species. Tissue-specific conditional knockout and rescue experiments reveal muscle as a critical site of dMtx2 action, with alternations of myofibril assembly and myogenic proteins being observed in *dMtx2* mutants. Structural and functional mitochondrial abnormalities are also detected, verifying dMtx2’s function in mitochondrial homeostasis. Notably, Mtx2 deficiency affects beta-barrel protein biogenesis and muscle development in pupa but not in larva, demonstrating Mtx2’s dynamic regulation in mitochondrial proteostasis and muscle development. Our results elucidate mitochondrial mechanisms driving potential muscle-autonomous defects in MADaM patients and highlights stage-specific Mtx2 function as a prospective therapeutic target for this progeroid syndrome.

## Introduction

The metaxin family comprises conserved mitochondria outer membrane (MOM) proteins encoded by nuclear genes, with three members (Mtx1, Mtx2, and Mtx3) identified in vertebrates (1). Mtx1, the first characterized member, was initially discovered in human due to its genomic proximity to *thrombospondin 3* (TSP 3) and *glucocerebrosidase* (GC) (2). Structurally, it contains two glutathione S-transferase (GST) domains and a C-terminal mitochondrial targeting sequence (MTS) that directs its subcellular localization (3, 4). Mtx2, discovered as an Mtx1-interacting protein through yeast two-hybrid screens, shares similar structural homology with Mtx2 but lacks an MTS (5). In contrast, Mtx3, identified in zebrafish and frogs, remains poorly characterized, with its functional role yet to be elucidated (1, 6).

Metaxins are evolutionarily conserved orthologs of yeast Sam37 (TOM37), a component of the sorting and assembly machinery (SAM) complex (7–9). In animals, the SAM complex— comprising Mtx1, Mtx2, and Sam50—orchestrates the biogenesis of mitochondrial β-barrel proteins, including Sam50 itself, the metabolic channel VDAC (Voltage-dependent anion channel), and Tom40, the central pore of the translocase of the outer membrane (TOM) complex (10). These β-barrel proteins are indispensable for mitochondrial physiology: Sam50 anchors the SAM complex (11), VDAC facilitates metabolite exchange across the MOM (12), and Tom40 mediates the import of nuclear-encoded mitochondrial pre-proteins (13). Perturbation in SAM complex components impairs β-barrel protein assembly, compromising mitochondrial function and cell viability across species (14, 15).

Beyond protein biogenesis, the SAM complex influences mitochondrial architecture and cellular signaling. Sam50 interacts with the MICOS (mitochondrial contact site and cristae organizing system) complex subunit Mic19, linking SAM to cristae maintenance; depletion of Sam50 or Mtx2 disrupts cristae morphology (16, 17). Metaxins also regulate extrinsic apoptosis by modulating Bak activation during TNFα signaling (18–20), and promote mitochondria fission via recruitment of the actin motor protein Myo19 (21).

Human pathologies underscore the importance of metaxins. Recent studies link *Mtx2* mutations to a new type of mandibuloacral dysplasia (MADaM), a progeroid syndrome marked by growth retardation, craniofacial anomalies, and musculoskeletal and skin malformations (22–25). MADaM-associated *Mtx2* mutations include splice-altering indels and missense mutations that are predicted to produce truncations with abolished protein function (22). Fibroblasts from MADaM patients exhibit mitochondrial fragmentation, respiratory chain deficits, TNFα resistance, verifying the known function of Mtx2 in mitochondrial homeostasis (22). Additionally, nuclear abnormalities and premature cellular senescence were also identified in patient fibroblasts, suggesting shared pathogenic mechanisms with classical MAD subtypes that are primarily caused by nuclear envelop protein defects (e.g., *LMNA* and *ZMPSTE24*) (26).

Animal models have been instrumental in dissecting Mtx2’s pathophysiology. Global *Mtx2* knockout causes early embryonic lethality in mice (17), while podocyte-specific deletion causes glomerular defects, partially recapitulating kidney manifestations in MADaM patients (24). Liver-specific Mtx2 ablation in mouse activates cGAC-STING-mediated IFN-I response and induces sterile inflammation (27). *C. elegans Mtx-2* mutants partially mirror MADaM phenotypes, exhibiting mitochondrial defects, developmental delay, and cuticle abnormalities (22, 28). Neuronal Mtx2 in flies and worms associates with kinesin/dynein motors to regulate mitochondria transport (29, 30).

Despite these advances, key questions persist. Although Mtx2 is ubiquitously expressed, MADaM manifests tissue-specific vulnerabilities, yet the direct versus systemic contributions of Mtx2 in affected tissues remain unclear. For example, it remains unknown what roles dMtx2 has in musculoskeletal system and what cause the growth retardation and myopathy in MADaM patients. Developmental stage-specific requirements of Mtx2 for beta-barrel protein biogenesis and mitochondrial physiology is key for explaining why progeroid symptoms emerge at certain developmental stage, but remain unexplored. Existing models also face limitations: *C. elegans* Mtx-2 is not functionally complemented by human orthologs, and murine early embryonic lethality precludes detailed developmental studies. Thus, novel models are needed to unravel tissue- and stage-specific pathophysiology and enable therapeutic discovery.

Here, we investigate *Drosophila* as a novel model for studying the developmental and tissue-specific functions of Mtx2. *Drosophila Mtx2* (*dMtx2*) null mutants exhibit recessive late pupal lethality, rescued by transgenic expression of either *Drosophila* or human *Mtx2*. Tissue-specific analyses pinpoint muscle as a critical site of dMtx2 function, with loss of dMtx2 arresting myofibrillogenesis, disrupting mitochondrial ultrastructure and ATP production, and downregulating mitochondrial and myogenic proteins. Strikingly, while dMtx2 is essential for Tom40 biogenesis in pupal muscles, larval muscles tolerate its loss, implicating developmental regulation of β-barrel assembly as a mechanism underlying tissue-specific vulnerabilities. Our findings establish *Drosophila* as a powerful model to dissect Mtx2’s roles in mitochondrial proteostasis, muscle development, and MADaM pathogenesis, offering new avenues for mechanistic and therapeutic exploration.

## Results

### Metaxin-2 deficiency causes *Drosophila* pupal lethality

We previously showed that dMtx2 deficiency results in complete pre-adult lethality using two null alleles, *dMtx2^1^* and *dMtx2^ΔDsRed^* (Fig. 1A) (30). To further investigate this lethal phenotype, we examined animal survival at multiple developmental stages and found that *dMtx2* null mutants survived normally during larval and pupal stages, with no obvious difference in larval body length compared to control animals (Fig. 1B and 1C). The *dMtx2* mutant animals exhibited no defects in external morphologic transitions until late pupal stages (Fig. 1D and S1). At 4 days after pupal formation (4d APF or P4d), a mild growth delay was observed, characterized by lighter epidermal pigmentation of the mutant pupae compared to control animals (Fig. 1D). Wild type adults normally emerge at 4.5 days after pupation, while mutant pupae failed to eclose and survived prolonged pupal stages for up to P7d with apparent heartbeats.

**Fig. 1.**
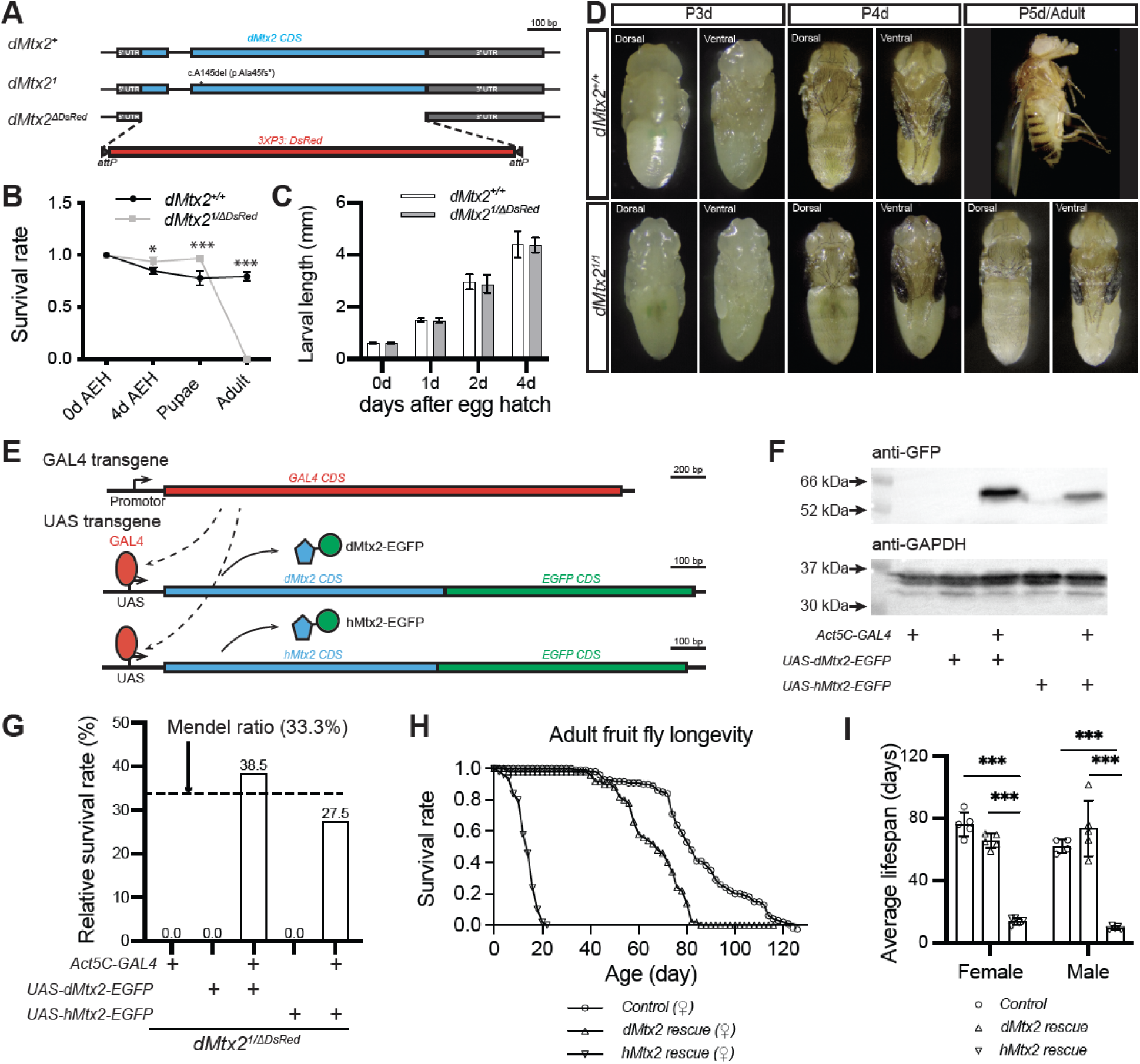
Metaxin-2 (Mtx2) is essential for pre-adult viability in Drosophila. **(A)** Schematic diagrams of the *Drosophila Mtx2* gene locus and two null mutations. **(B)** Survival rates of control and *dMtx2^1/ΔDsRed^* mutant animals across developmental stages (two-way ANOVA, Bonferroni test; *N*=3 tests). AEH: after egg hatch. **(C)** Body length measurements of control and *dMtx2* mutant larvae (two-way ANOVA, Bonferroni test; *N*=19∼75 animals). **(D)** Morphologic comparison of control and mutant pupae/adult at 3-5 days after pupae formation. **(E)** Schematic of GAL4/UAS-driven *dMtx2-EGFP* and *hMtx2-EGFP* transgenes. **(F)** Western blot of adult lysates showing transgenic dMtx2-EGFP and hMtx2-EGFP expression. **(G-I)** Rescue of *dMtx2^1/ΔDsRed^* lethality by *dMtx2-EGFP* and *hMtx2-EGFP* transgenes using ubiquitous *Act5c-GAL4*. Relative survival rate quantifies the rescue effect of pre-adult lethality (G). The survival rates for each genotype are listed above the bars. Survival curves (H) and average lifespans (I) of control and rescued adults demonstrate the long-term rescue effects. Data in panel I were analyzed by two-way ANOVA, Bonferroni test; N=4∼5 tests. Statistical significance in all figures: * *P*<0.05; ** *P*<0.01; *** *P*<0.001. Error bars in all figures are standard deviation of the mean.

To further confirm dMtx2’s essential role in animal survival, we generated a UAS transgene to express EGFP-fused dMtx2 through the GAL/UAS system (Fig. 1E and 1F) (31). When dMtx2-EGFP was ubiquitously expressed by *Act5C-GAL4* in the mutant flies, the animals were able to survive to adulthood (Fig. 1G) and had normal lifespan as the control flies (Fig. 1H and 1I). Using this rescue paradigm, we examined the functional similarity between *Drosophila* and human Mtx2. A *UAS-hMtx2-EGFP* transgene was generated in the same way as the *UAS-dMtx2-EGFP* strain (Fig. 1E and 1F). When hMtx2-EGFP was expressed in *dMtx2* mutant flies, the pupal lethality was largely rescued, albeit with milder efficacy than dMtx2-EGFP (Fig. 1G). The hMtx2-rescued flies displayed a much shorter lifespan than control or dMtx2 rescued flies (Fig. 1H and 1I), demonstrating a partial compensation of hMtx2 for dMtx2 in *Drosophila*. These results suggest conserved molecular functions between hMtx2 and dMtx2, despite of a low sequence identity shared by these two proteins (Fig. S2A)

MADaM-associated *hMtx2* mutations are predicted to produce truncated proteins (Fig. S2B). However, whether these mutations are truly dysfunctional have not been experimentally tested. We generated transgenes to express three pathogenic *hMtx2* mutations in *Drosophila* and conducted rescue experiments to assess their functions. In contrast to the wildtype *hMtx2*, none of the mutated *hMtx2* transgene was able to rescue the pupal lethality of *dMtx2* null mutants (Fig. S2C), verifying the loss-of-function (LOF) nature of these mutations.

### Muscle dMtx2 is indispensable for animal survival

Given that *dMtx2* mutants die in late pupal stages without apparent morphologic defects, we wondered which cell/tissue types require dMtx2 for animal survival. Using an EGFP knock-in allele, *dMtx2^EGFP^*, we have shown that dMtx2 is broadly expressed in *Drosophila* tissues and in multiple developmental stages (30). The *dMtx2-EGFP* coding sequencing could be excised out through flippase (FLP)-mediated FRT recombination so that *dMtx2^EGFP^*also serves as a conditional knockout (cKO) allele (Fig. 2A). For example, dMtx2-EGFP expression was efficiently abolished when *Tubulin-GAL4* was utilized to express FLP ubiquitously (Fig. 2B and 2C). This global knockout resulted in late pupal lethality, similar to *dMtx2* null mutants. The lethality was completely rescued by co-expression of the *UAS-dMtx2-EGFP* transgene, confirming the specificity of the *dMtx2* cKO (Fig 2D).

**Fig. 2.**
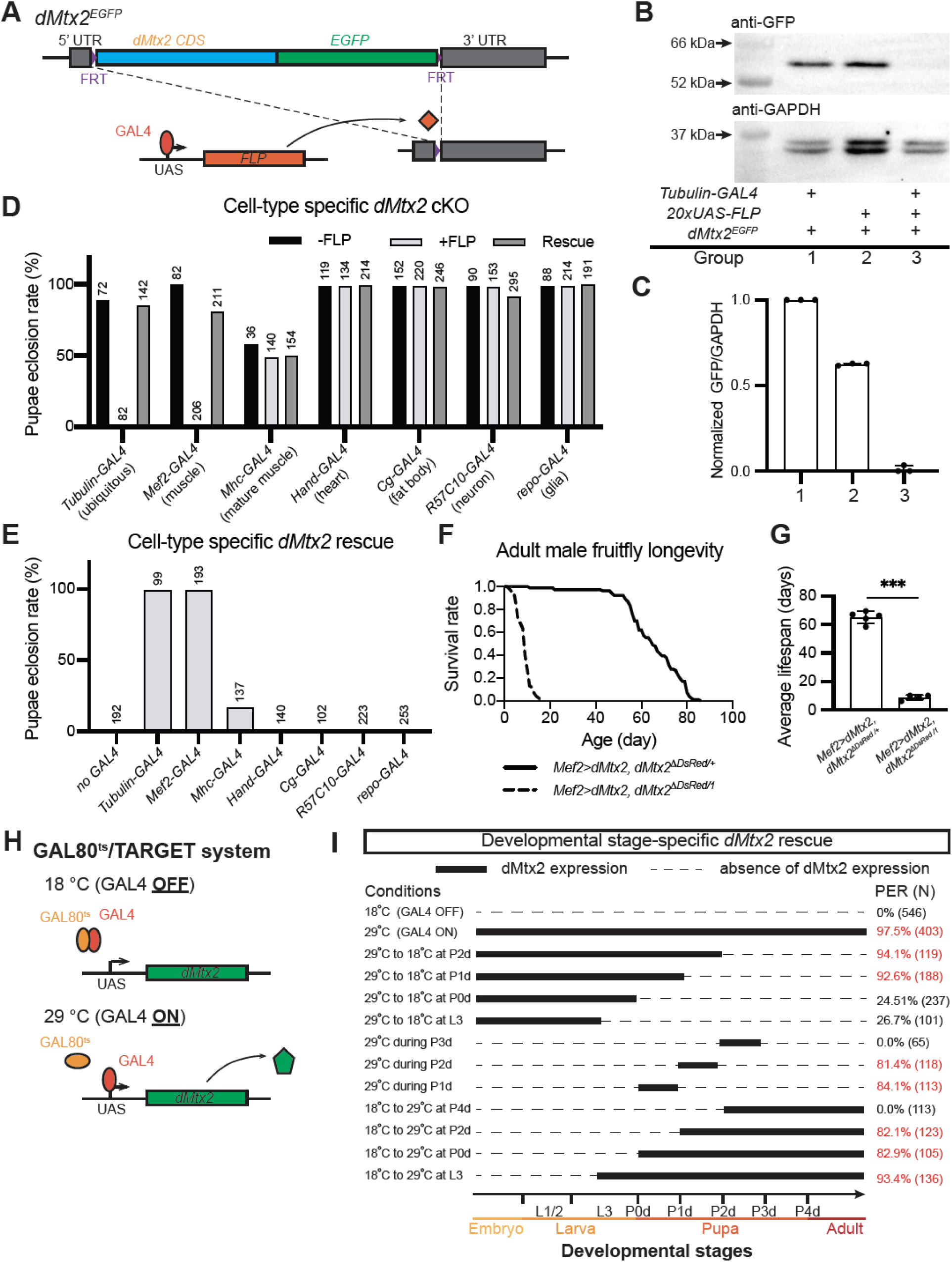
Muscle-specific dMtx2 is essential for viability. **(A)** Strategy for FLP/FRT-mediated conditional knockout (cKO) using *dMtx2^EGFP^*. **(B-C)** Western blot (B) and quantification (C) of dMtx2-EGFP expression in control (without *tubulin-GAL4* or *20xUAS-FLP*) or cKO (*w; tubulin-GAL4/+; dMtx2^EGFP^, 20xUAS-FLP/+*) larvae (Kolmogorov-Smirnov test; *N*=3 tests). **(D)** Ubiquitous and tissue-specific GAL4s were used to express FLP in *dMtx2^1/EGFP^* flies. Control flies did not have *UAS-FLP* and rescue flies had *UAS-dMtx2-EGFP*. Pupae eclosion rates (PERs) were quantified to evaluate the cKO effects. **(E)** Ubiquitous and tissue-specific GAL4s were used to express transgenic *dMtx2-EGFP* in *dMtx2^1/ΔDsRed^* flies and PERs were quantified to evaluate the rescue effects. The number of animals used for each test are listed above the bars. **(F-G)** Survival curves (F) and average lifespans (G) of control (*w; UAS-dMtx2-EGFP/+; Mef2-GAL4, dMtx2^ΔDsRed^/dMtx2^+^*) and muscle-rescued (*w; UAS-dMtx2-EGFP/+; Mef2-GAL4, dMtx2^ΔDsRed^/dMtx2^1^*) adult flies (Welch’s t test, *N*=4∼5 tests). **(H)** Schematic of TARGET system using temperature-sensitive GAL80 (GAL80^ts^). **(I)** *Tub-GAL80^ts^*was used to control *Mef2-GAL4* activation and *UAS-dMtx2-EGFP* expression in *dMtx2* null mutant flies (*w; UAS-dMtx2-EGFP/tub-GAL80^ts^; Mef2-GAL4, dMtx2^ΔDsRed^/dMtx2^1^*) in developmental stage-specific fashions by raising animals at different temperatures.

We then conducted cell-type specific KO experiments with multiple GAL4 drivers, including *Mef2-GAL4* in myoblasts and muscles, *Mhc-GAL4* in differentiated muscles, *Hand-GAL4* in hearts, *Cg-GAL4* in fat bodies, *R57C10-GAL4* in neurons and *repo-GAL4* in glial cells. Among these drivers, only *Mef2-GAL4*‒mediated dMtx2 knockout gave complete pupal lethality (Fig. 2D), indicating that muscle dMtx2 is required for animal lethality. The Mef2-GAL4‒mediated lethality was fully rescued by co-expression of *UAS-dMtx2-EGFP* (Fig. 2D), confirming the muscle-autonomous function of dMtx2. Interestingly, *Mhc-GAL4*, which is expressed in differentiated muscles from middle pupal stages, failed to induce pupal lethality compared to controls, suggesting stage-dependent requirement of Mtx2 in muscles.

To further examine the importance of muscle Mtx2, we conducted cell-type specific rescue experiments by using different GAL4 drivers to express dMtx2 transgene in null mutants (Fig. 2E). Among all the drivers tested, ubiquitous *Tub-GAL4* and muscle-specific *Mef2-GAl4* were able to fully rescue mutant flies to adulthood, while mature muscle-specific *Mhc-GAL4* demonstrated a partial rescue effect (Fig. 2E). In summary, the cKO and rescue results suggest muscle dMtx2 expression is required and sufficient for supporting pre-adult survival. However, unlike the *Act5C-GAL4* rescue flies (Fig. 1H and 1I), the *Mef2-GAL4* rescue flies had a much shorter lifespan (Fig. 2F and 2G), indicating that dMtx2 expression in other tissues is also important for adult health and survival.

### Muscle dMtx2 is required for animal survival in a developmental stage-dependent manner

The distinct outcomes of *Mef2-GAL4* and *Mhc-GAL4* in cKO and rescue experiments suggests that dMtx2 exerts stage-dependent functions in muscles. To determine the developmental time window requiring dMtx2 activity, we performed stage-specific dMtx2 rescue experiments using the TARGET system, which employs temperature-sensitive *Tub-GAL80^ts^* to temporally control GAL4 activity and UAS transgene expression. In this system, GAL80^ts^ suppresses GAL4 activity at permissive temperature (18 °C), but lost its inhibitory ability at restrictive temperature (29 °C) (Fig. 2H).

To validate the TARGET system’s efficacy, we firstly tested muscle-specific dMtx2 rescue under constant temperatures. At 18 °C (GAL80 active/GAL4 suppressed), all animals (*N*=546) died during late pupal stages, whereas at 29 °C (GAL80 inactive/GAL4 active), most pupae (97.5%, *N*=403) successfully eclosed (Fig. 2I), confirming temperature-dependent control of transgene expression.

Next, we mapped the critical developmental window by shifting animals between 18 °C and 29 °C to activate *Mef2-GAL4>dMtx2* at specific stages (Fig. 2I). GAL4 activation that was initiated from the early embryonic stages but terminated prior to puparium resulted in partial rescue (20∼30% PER). GAL4 activation that was initiated after the second puparium day (>P2d) failed to restore viability (0% PER). In contrast, the paradigms with GAL4 activation during the early pupal phase (first two days after pupal formation, P1d–P2d) restored robust eclosion (PER >80%), with even transient expression limited to P1d or P2d alone rescuing viability. Notably, P2d emerged as the pivotal stage: *dMtx2* mRNA and protein produced during P1d could persist into P2d, suggesting that sustained protein levels during P2d are essential. These results identify a narrow developmental window—centered on the second puparium day (P2d)—during which dMtx2 is indispensable for pupal eclosion.

### Loss of dMtx2 causes pupal muscle growth defects

To investigate dMtx2’s functions in muscles, we examined the growth of dorsal longitudinal muscles (DLMs), a group of indirect flight muscles responsible for the fast beating of fly wings, during pupal stages. We stained actin filament and integrin beta subunit (βPS) to highlight muscle morphologies and muscle-cuticle attachments (Fig. 3A and S3A). The number of DLMs, the general muscle arrangement patterns, and muscle-cuticle junction structures appeared similar between *dMtx2^1^* heterozygous and homozygous animals (Fig. 3A), suggesting that early muscle development events—such as myoblast proliferation and fusion, and myotube targeting and attachment—are largely unaffected in *dMtx2* mutants. However, muscle fiber sizes differed between mutant and control animals, with mutant muscles becoming significantly smaller in late pupal stages (Fig. 3A and S3A). At P4d, control DLMs grew to full capacity and occupied the entire thoracic space. In contrast, large gaps remained between muscle fibers in mutant animals. Muscle size difference was validated by quantifying DLM4 size in sagittal sections (Fig. 3B). Since *dMtx2* mutant animals survived additional days within the pupal case, we wondered whether mutant muscles continued to grow after P4d. The quantification of DLM4 size showed similar muscle sizes during the extended pupal period (Fig. S3B and S3C), suggesting that dMtx2 deficiency halted, rather than delayed muscle growth in late pupal stages. Furthermore, co-expression of the dMtx2 transgene by *Mef2-GAL4* rescued the muscle size reduction (Fig. 3A and 3B), suggesting a cell type-autonomous function of dMtx2 in muscle growth.

**Fig. 3.**
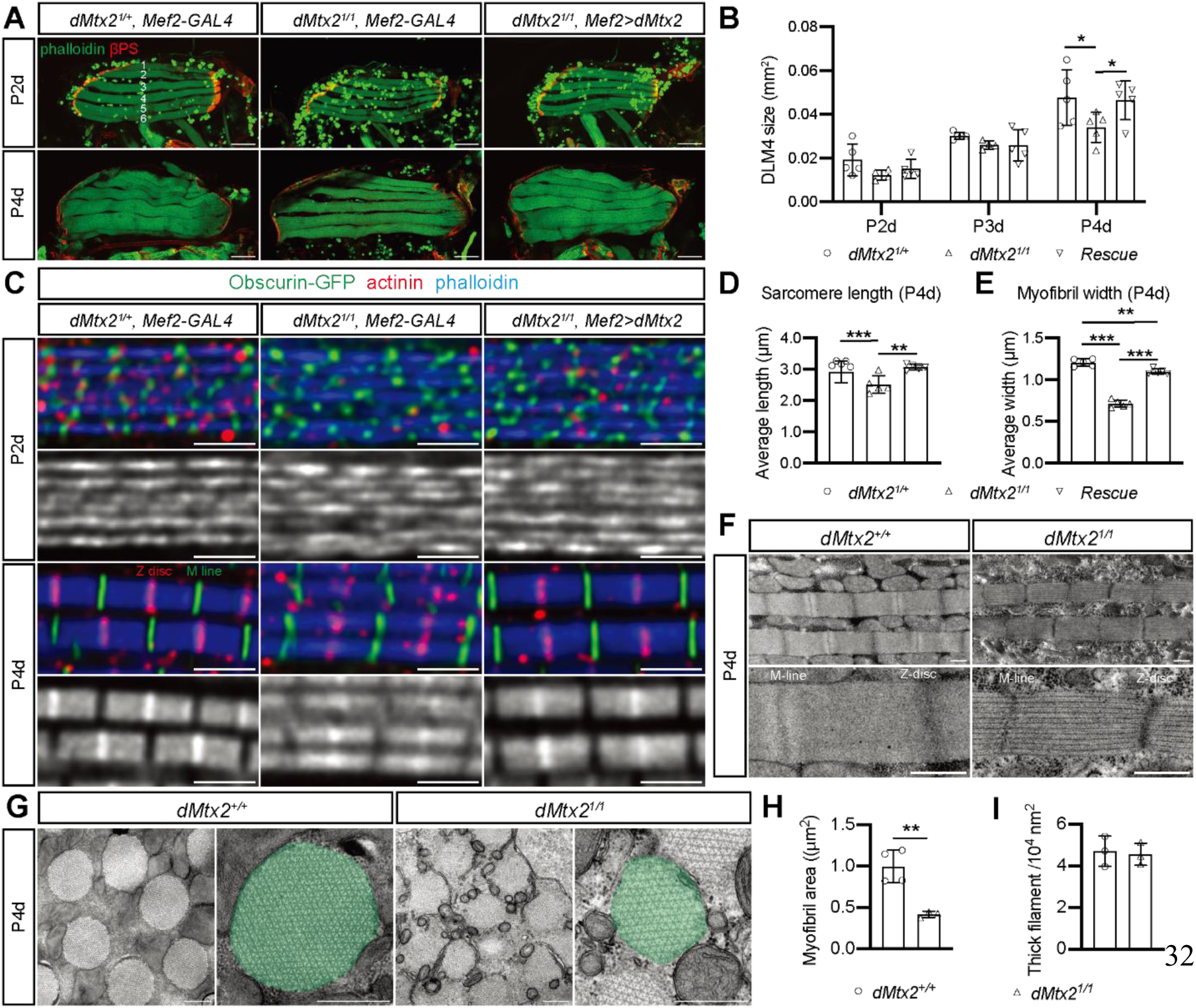
dMtx2 loss disrupts pupal muscle growth. **(A)** Phalloidin (F-actin, green) and βPS-integrin (red) staining of dorsal longitudinal muscles (DLMs) in pupae. Scale: 100 µm. **(B)** DLM4 size quantification (two-way ANOVA, Tukey’s test; *N*=5 animals). **(C)** Sarcomere structures labeled with phalloidin (blue), Obscurin-GFP (M-line, green), and actinin (Z-disc, red) at 2 or 4 days after pupal formation. Scale: 2 µm. **(D-E)** Measurements of sarcomere length (D) and myofibril width (E) in late pupae (one-way ANOVA; *N*=5 animals). **(F–G)** TEM of longitudinal (F) and cross-sectional (G) DLMs. Scale: 500 nm. **(H–I)** Measurements of myofibril area (H) and thick filament density (I) on TEM micrographs (Welch’s t test; *N*=3∼4 animals).

Beyond muscle morphology, immunohistochemistry was performed to reveal the details of muscle growth defects in *dMtx2* mutants. Sarcomere substructures were labeled with phalloidin for thin filaments, actinin or kettin for Z-disc, and Obscurin-GFP for M-line (Fig. 3C and S3D). At the middle pupal stage (P2d), myofibrils are thin and sarcomeres just begin to form. Obscurin-GFP is already localized to the M-line but actinin puncta are not restricted to Z-discs. During late pupal stages, myofibrils became much thicker, while actinin/kettin and Obscurin-GFP formed sharp lines at Z-discs and M-lines, respectively, indicating sarcomere maturation. In mutant muscles, while initial sarcomere formation was similar to controls at P2d, late-stage pupae exhibited thinner myofibrils and shorter sarcomeres (Fig. 3C-E, S3D). The Z-disc and M-line markers also became less organized in *dMtx2* mutant muscles, together showing sarcomere maturation defects. Again, these phenotypes were successfully rescued when transgenic dMtx2-EGFP was co-expressed in the mutant muscles (Fig. 3C-E, S3D).

Transmission electronic microscopy (TEM) further illustrated ultrastructural changes in flight muscles from late-stage pupae. Smaller sarcomeres were found in longitudinal sections, with differences in electron density, especially in the M-line, suggesting altered composition and construction of sarcomeric structures (Fig. 3F). Changes in myofibril diameter were more apparent in cross-sections (Fig. 3G), with an average myofibril area of 0.996 µm^2^ in control muscles and 0.418 µm^2^ in mutant muscles (Fig. 3H). In contrast, the general organization and density of thick filaments remained unchanged in the mutant (Fig. 3I).

### Mitochondrial structure and function are compromised in *dMtx2* mutant

By using the *dMtx2^EGFP^* allele, we showed clearly that dMtx2 is located to the mitochondrial outer membrane in *Drosophila* muscles (Fig. 4A). Thus, loss of dMtx2 might primarily cause mitochondrial defects and subsequently affect muscle development. In TEM sections of fight muscles, smaller mitochondria were much more abundant in *dMtx2* mutant muscles compared to the controls (Fig. 4B), with the average mitochondrial cross-section area reduced from 0.308 µm^2^ in controls to 0.054 µm^2^ in mutant animals (Fig. 4C).

**Fig. 4.**
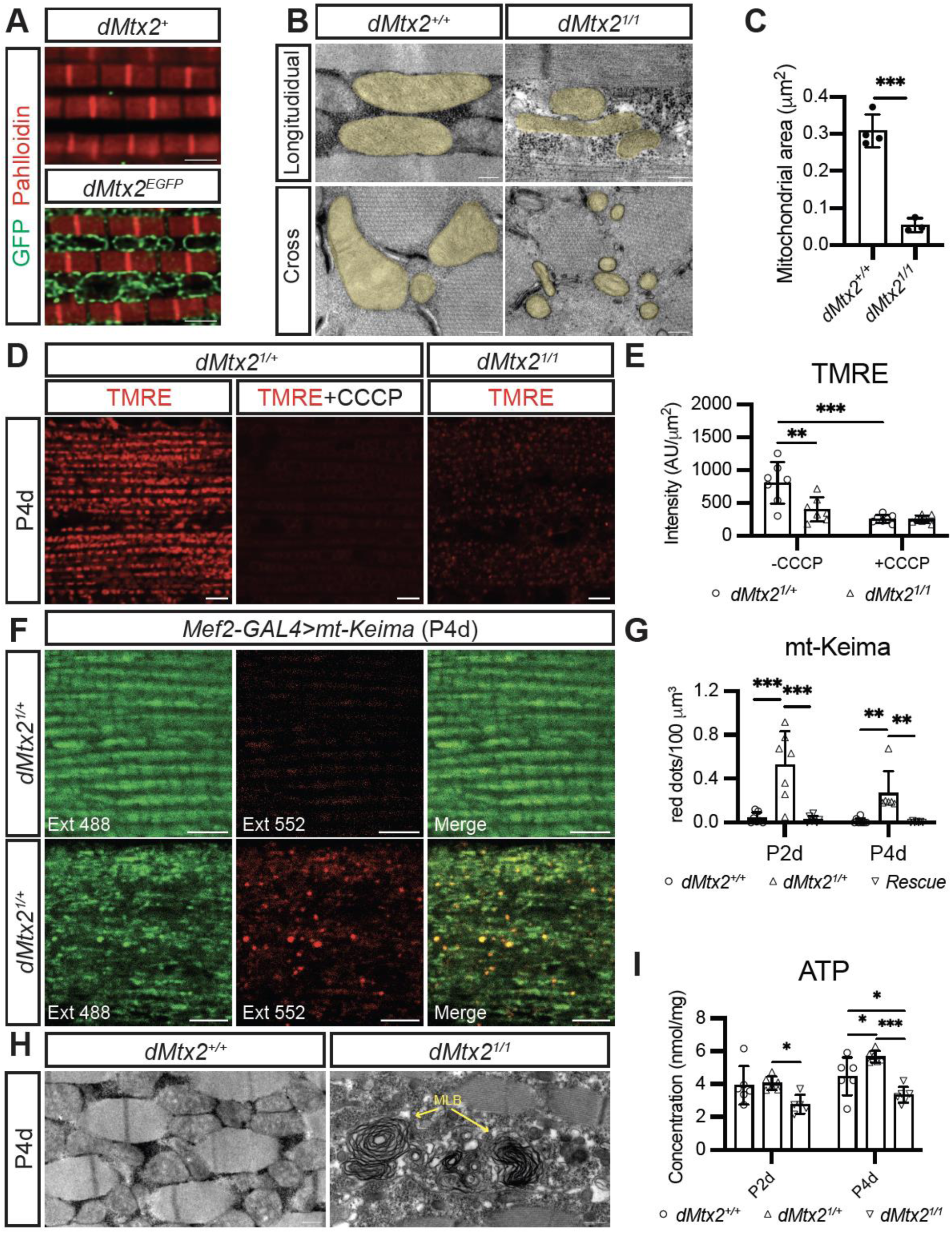
dMtx2 deficiency causes mitochondrial structural and functional defects in pupal muscles. **(A)** Endogenous dMtx2-EGFP (green) and phalloidin staining (red) in adult DLMs. Scale bars: 2 µm. **(B)** TEM of mitochondria in P4d DLMs. Scale bars: 200 nm. (C) Mitochondrial size quantification in TEM cross sections (Welch’s t test; *N*=3∼4 animals). **(D-E)** TMRE fluorescence (D) and quantification (E) in pupal DLMs for testing mitochondrial membrane potential (ΔΨm) (Two-way ANOVA, Turkey’s test; *N*=7 animals). Scale bars: 5 µm. **(F-G)** mt-Keima fluorescence and quantification in pupal DLMs for detecting mitophagy (Two-way ANOVA, Turkey’s test; *N*=6∼7 animals). Scale bars: 5 µm. **(H)** Multilayered bodies (MLB) (yellow arrows) in TEM images. Scale bars: 500 nm. **(I)** ATP levels in whole pupal lysates (Two-way ANOVA, Turkey’s test; *N*=5∼6 tests).

In addition to the mitochondrial structural changes, mitochondrial function was also assessed. We used TMRE (tetramethylrhodamine ethyl ester) to detect mitochondrial membrane potential (ΔΨm) (Fig. 4D and 4E). Control muscles exhibited high-intensity TMRE fluorescence, which was effectively abolished by CCCP (carbonyl cyanide 3-chlorophenylhydrazone) treatment, confirming successful mitochondrial incorporation of the probe. Strikingly, dMtx2 mutant displayed a prominent reduction in TMRE fluorescence intensity compared to controls, demonstrating a significant decrease in ΔΨm.

Mitochondrial depolarization often activates mitophagy signaling pathway and induces the formation of autolysosome. To detect mitophagy, we employed the *Mef2-GAL4* driver to express *UAS-mt-Keima* in muscles. In control DLMs, mt-Keima was excited by a 488-nm laser but rarely by a 552-nm laser (Fig. 4F), indicating that mitochondria were in a neutral pH environment. In contrast, the number of 552-nm‒excited puncta were significantly increased in mutant DLMs at both early and late pupal stages (Fig. 4F and 4G), indicating an increase in mitochondria acidification. The formation of autophagosomes were also verified by TEM data, with multilamellar bodies (MLBs) detected in mutant muscles but rarely in controls (Fig. 4H). Our results show that mitophagy and autophagy are up-regulated in *dMtx2* mutant muscles at P2d suggesting that mito-lysosome formation begins at the onset of myofibrillogenesis, prior to the sarcomere maturation defects observed in *dMtx2* mutant muscles.

Furthermore, we measured ATP concentration in whole pupal tissues to address whether mitochondrial defects impact energy production. Control animals exhibited progressive elevation of ATP levels from early to late pupal stages, demonstrating enhanced mitochondrial activities during pupal development (Fig. 4I). However, mutant animals displayed significantly reduced ATP concentrations at both pupal stages, reflecting impaired mitochondrial functionality. In summary, we identified various mitochondrial defects in *dMtx2* mutant animals that occur prior to the emergence of myofibril alternations and likely have a strong influence on muscle development.

### Loss of dMtx2 reduces *Drosophila* Mtx1 and beta-barrel protein expression

Given that Mtx2 is post-translationally required for Mtx1 and beta-barrel protein expression in yeast and mammalian cells, we examined the expression of CG9393 (Mtx1 homolog in *Drosophila*), Porin (*Drosophila* VDAC) and Tom40. Since no antibody against CG9393 was available, we generated a *CG9393* knock in allele (*CG9393^Myc^*) and successfully detected Myc-tagged CG9393 by Western Blot (Fig. 5A). To examine the function of dMtx2 on CG9393 expression, we employed the CRISPR-Cas9 technique to globally knock out dMtx2, and observed a concurrent depletion of Myc-tagged CG9393 in pupal tissues (Fig. 5A and 5B), suggesting that Mtx1/2 interactions are conserved in *Drosophila*.

**Fig. 5.**
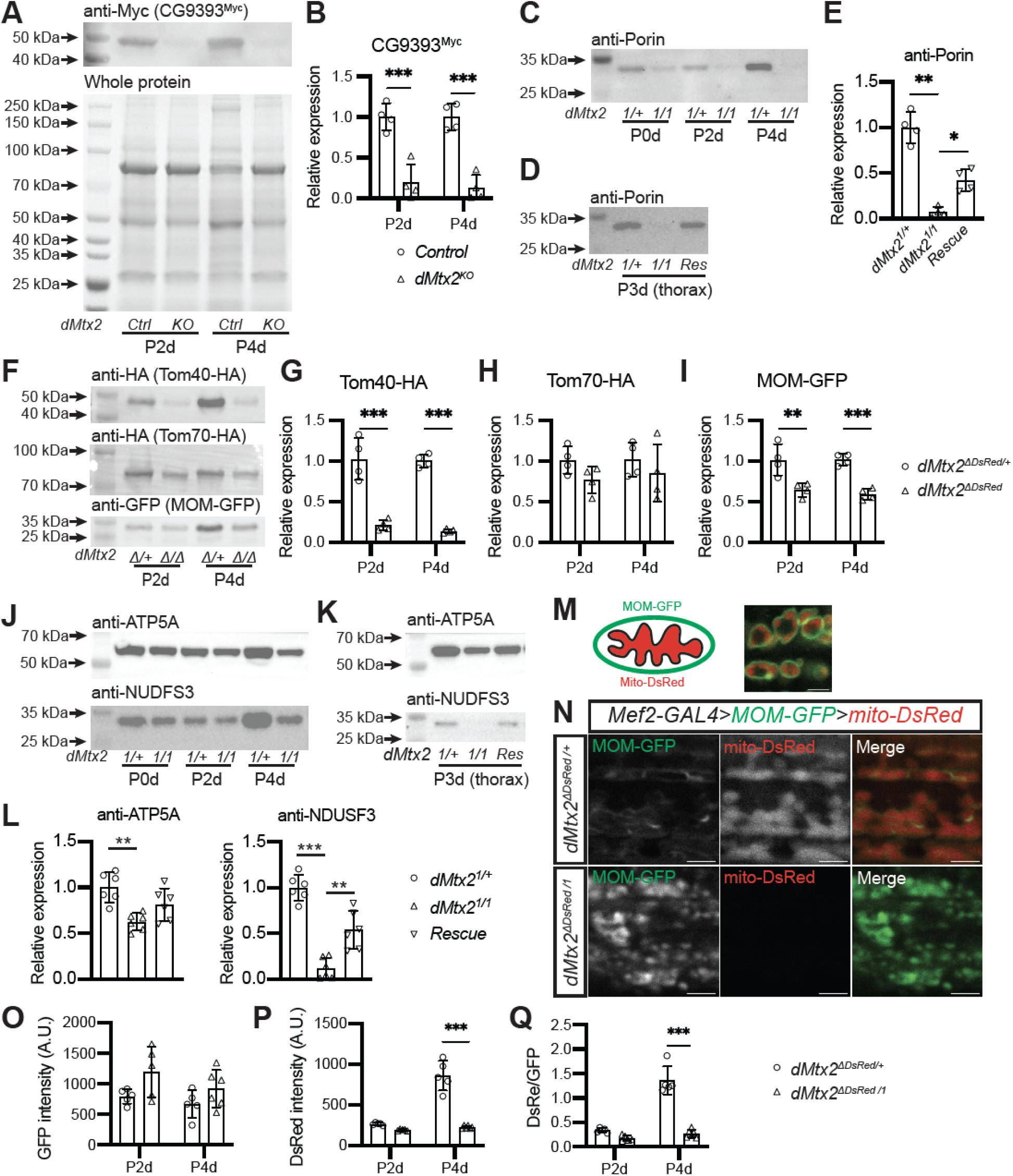
Loss of dMtx2 compromises mitochondrial protein expression and import. **(A-B)** Western blot (A) and quantification (B) of CG9393-Myc expression in control (Ctrl, *w; Tubulin-GAL4/UAS-Cas9; CG9393^Myc^/+*) or dMtx2 knockout (KO, *w; Tubulin-GAL4/UAS-Cas9, U6:dMtx2^sgRNA^; CG9393^Myc^/+*) pupae. Whole protein quantity is analyzed by using the Stain-free SDS-PAGE (Bio-Rad) and is used as loading controls. All values were normalized to the mean value of the control group (Two-way ANOVA, Bonferroni test; *N*=4 tests). **(C)** Porin expression across pupal stages by Western blot. **(D–E)** Western blot (D) and quantification (E) of Porin expression in P3d thorax lysates of *dMtx2* heterozygous, homozygous and muscle-rescue homozygous animals (Brown-Forsythe and Welch ANOVA test; *N*=4 tests). **(F–I)** Western blots (F) and quantifications of transgenic Tom40-HA (G), Tom70-HA (H), and MOM-GFP (I) expression in pupal muscles using *Mef2-GAL4* (Two-way ANOVA, Bonferroni test; *N*=4 tests). **(J–L)** ATP5A and NDUFS3 expression by Western blot (Brown-Forsythe and Welch ANOVA test; *N*=6 tests). **(M)** Schematic and a fluorescent image of MOM-GFP and mito-DsRed localization in mitochondria. **(N-Q)** Fluorescent images (N) and quantifications (O-Q) of MOM-GFP and Mito-DsRed in P4d DLMs (Two-way ANOVA, Bonferroni test; *N*=5∼6 animals).

To investigate how dMtx2 regulates beta-barrel proteins, an anti-Porin antibody was used to examine Porin expression in multiple pupal tissues by Western blot (32). Compared to controls, reduced Porin expression was detected in mutant animal, with the reduction becoming more prominent in later pupal stages (Fig. 5C). Muscle-specific expression of the *dMtx2* transgene significantly restored Porin levels in pupal thoracic tissues (Fig. 5D and 5E). To validate the expression of another beta-barrel protein Tom40, we expressed transgenic Tom40-HA in muscles using *Mef2-GAL4* and detected its expression using an anti-HA antibody (Fig. 5F). Tom40-HA expression was reduced by more than 80% on average in *dMtx2* mutant muscles compared to control muscles (Fig. 5F and 5G). Co-expression of dMtx2-EGFP successfully rescued the loss of Tom40-HA in *dMtx2* mutant muscles (Fig. S4A and S4B).

For comparative analysis, we also examined the expression of two alpha-helical MOM proteins, Tom70-HA and MOM-GFP. Tom70 is a peripheral receptor of TOM complex (33), and MOM-GFP is a non-functional fusion protein composed of rat Tom20 mitochondrial targeting sequence and EGFP (34). In contrast to Porin and Tom40-HA, the expression of Tom70-HA and MOM-GFP was only mildly affected in *dMtx2* mutant pupae (Fig. 5F-I), showing the specific regulation of *dMtx2* on the expression of β-barrel proteins in mitochondrial outermembrane.

### Loss of dMtx2 impairs mitochondrial protein import

Given the essential role of Tom40 in TOM complex, dMtx2 deficiency is expected to impact mitochondrial protein import. As examples, we examined the expression of ATP5A and NUDFS3, which are Complex I and Complex V components in the mitochondrial inner membrane (MIM). Western blot analyses showed that both proteins were gradually downregulated in *dMtx2* mutant pupae, with NUDFS3 being more affected than ATP5A (Fig. 5J). Similar to Porin and Tom40, the expression of both MIM proteins was partially restored by co-expression of the dMtx2 transgene in pupal muscles (Fig. 5K and 5L).

To further assess Mtx2’s function in mitochondrial protein import, we acquired additional evidence using MOM-GFP and mito-DsRed, a mitochondrial matrix reporter created by fusing human COX8A minimal MTS with DsRed (35). When these two proteins were co-expressed in fly muscles, high-resolution fluorescent confocal imaging clearly distinguished their distinct localizations in mitochondrial outer membrane and matrix (Fig. 5M). We then compared their fluorescent intensities in control and *dMtx2* mutant muscles. With MOM-GFP, we observed a clear change of mitochondrial morphology in mutant muscles, with MOM-GFP itself becoming more condensed and the average fluorescence intensity of MOM-GFP being slightly increased in mutant tissues (Fig. 5N and 5O). In contrast, mito-DsRed fluorescence was greatly reduced in mutant animals, particularly at late pupal stages (Fig. 5N and 5P). The mito-DsRed to MOM-GFP fluorescence intensity ratio was significantly reduced in *dMtx2* mutant pupae (Fig. 5Q), suggesting that loss of dMtx2 could affect the localization of mitochondrial matrix proteins independent of its function, which is likely due to the disruption of mitochondrial protein import.

### Proteomic characterization reveals extensive alternations of mitochondrial protein expression in *dMtx2* mutant

Given the observed mitochondrial protein expression changes, we performed proteomic analyses to comprehensively characterize molecular alternations underlying mitochondrial and muscle developmental defects in *dMtx2* mutants. Since the second puparium day represents a critical time window for dMtx2-dependent development, we collected *dMtx2^1^* heterozygous and homozygous pupae at two days APF for tandem mass tag (TMT)-based quantitative proteomic profiling (Fig. 6A).

**Fig. 6.**
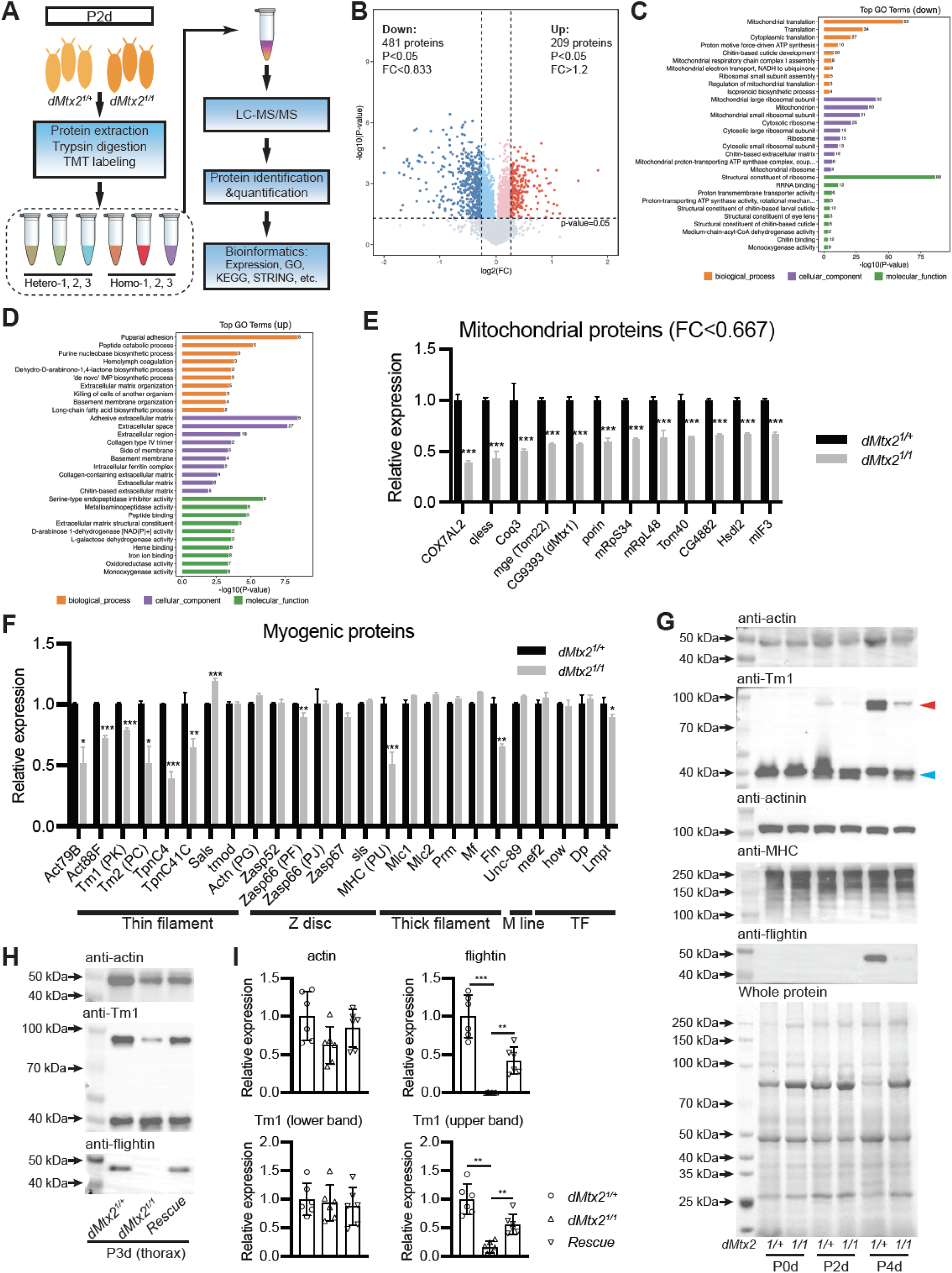
Proteomic analyses identified alternations of mitochondrial and muscle protein expression in *dMtx2* deficiency. **(A)** Workflow for pupal proteomics. **(B)** A volcano plot of differentially expressed proteins (DEPs) using low-stringency thresholds (|fold change(FC)|>1.2 or |log2(FC)|>0.263, *P*<0.05). **(C–D)** GO analyses of the downregulated (C) and upregulated (D) proteins. **(E)** A bar graph of the most impacted mitochondrial proteins ((|FC|>1.5) identified by MS/MS. **(F)** A bar graph of muscle protein levels by MS/MS. **(G)** Western blots of actin, tropomyosin-1 (Tm1), actinin, MHC and flightin (Fln) in whole animal lysates across pupal stages. Tm1 exhibits short and long isoforms (blue and red arrows). **(H-I)** Western blots (H) and quantification (I) of actin, Tm1 and Fln in P3d thorax lysates of *dMtx2* heterozygous, homozygous and muscle-rescue homozygous animals (Brown-Forsythe and Welch ANOVA test; *N*=6 tests).

Mass spectrometry detected nearly 6000 proteins in pupal lysate and over 5500 of them were qualified for statistical analyses (Data S1). Principal component analysis (PCA) revealed distinct expression profiles between heterozygous and homozygous groups (Fig. S5A). Using stringent statistical thresholds (|fold change(FC)|>1.2 or |log2(FC)|>0.263, *P*<0.05), we identified 690 differentially expressed proteins (DEPs), with 481 proteins downregulated and 209 proteins upregulated in homozygous mutants (Fig. 6B). Geno Ontology (GO) and Kyoto Encyclopedia of Genes and Genomes (KEGG) analyses of DEPs highlighted functional enrichment groups. Among the top enriched categories of downregulated DEPs included over 80 mitochondrial proteins, with 53 of them involved in mitochondrial ribosomal formation/translation and 21 in oxidative phosphorylation (Fig. 6C and S5B). The top enriched categories of up-regulated DEPs included adhesive extracellular proteins and metabolic proteins (Fig. 6D and S5C), which might reflect the developmental changes of *dMtx2* mutants.

To identify the most impacted mitochondrial proteins, we reapplied stricter thresholds (|fold change(FC)|>1.5 or |log2(FC)|>0.585, *P*<0.05), yielding 145 DEPs (Fig. S5D). Twelve downregulated mitochondrial proteins were identified, including CG9393, Porin and Tom40. Meg (*Drosophila* Tom22) was also significantly reduced (Fig. 6E). The variable reductions of Tom22 and other TOM complex components (Tom5, Tom6, Tom7, Tom20 and Tom70) were likely secondary to Tom40 loss (Fig. S6A). In addition to these MOM proteins, other most affected mitochondrial proteins included inner membrane and matrix proteins that are mostly involved in oxidative phosphorylation and mitochondrial translation (Fig. 6E). These results indicate widespread mitochondrial protein dysregulation, with TOM complex being directly and profoundly affected, in *dMtx2* mutants.

### Proteomic characterization reveals reduced expression of specific muscle proteins in *dMtx2* mutants

To identify muscle development-linked DEPs, STRING analysis of downregulated proteins from high-stringency criteria revealed an interacting subnetwork of muscle-related proteins, including actin, tropomyosin, troponin and myosin heavy chain (MHC) (Fig. S5E). Actin, the major component of thin filaments, is encoded by six *Drosophila* actin genes with stage- and tissue-specific expression (36). MS analysis detected four actins, two of which‒muscle-specific Act79B and Act88F‒were significantly reduced (Fig. 6F and S6B). Tropomyosins, coil-coil proteins critical for thin filament structure, are encoded by *Tm1* and *Tm2*, which produce multiple isoforms. Isoforms Tm1-PK and Tm2-PC were markedly decreased in mutants (Fig. 6F and S6C). Troponin, a ternary complex (TpnC, TpnH and TpnI), associating with actin filaments and regulating muscle contraction, includes TpnC isoforms generated by five tissue-specific genes (37). Adult muscle-specific TpnC4 and TpnC41C were significantly diminished in *dMtx2* mutants, while TpnH and TpnI (encoded by *up* and *wupA*, respectively) exhibited only mild isoform-specific reductions (Fig. 6F and S6D).

Myosin, the thick filament backbone, comprises two pairs of heavy chains, essential light chains, and regulatory light chains. *Drosophila* MHC is encoded by a single gene with multiple isoforms. MS detected four MHC isoforms, with only low-abundant isoform U reduced in *dMtx2* mutants (Fig. 6F and S6E). Myosin light chains (Mlc1, Mlc2) and structural protein paramyosin (Prm) and myofilin (Mf) remained unchanged. However, fight muscle-specific flightin (Fln) was significantly reduced (Fig. 6F). Z-disc and M-line scaffold proteins were largely unaffected, and key myogenic regulators (Mef2, how, Dp and Lmpt) showed no expression changes (Fig. 6F), suggesting that dMtx2 depletion selectively disrupts thin filament proteins without broadly altering transcriptional networks.

To validate the proteomic findings, we performed Western blot to assess muscle protein expression in pupal tissues (Fig. 6G). Total actin levels, derived from different genes with similar sequences and molecular weights, were mildly reduced in late pupal tissues (Fig. 6G). For Tm1, distinct long and short isoforms were differentially affected in *dMtx2* mutants: flight muscle-specific long isoforms (designated protein 33/34), which increased during pupal stages in controls, were selectively diminished in mutant tissues. While Z-disc protein actinin and a range of MHC isoforms were detected across multiple pupal stages, their expression remained unaltered in *dMtx2* mutants. In contrast, flightin, which accumulates in late pupae, exhibited significantly reduced expression in mutants. Co-expression of the *dMtx2* transgene driven by *Mef2-GAL4* (the ‘Rescue’ group) successfully restored Tm1(33/34) and Fln levels in pupal thorax (Fig. 6H and 6I), confirming that altered muscle protein expression arises cell-autonomously from *dMtx2* deficiency. Western blot results, consistent with MS data, underscore dMtx2’s role in regulating specific muscle proteins.

### dMtx2 is not essential for Tom40-dependent animal growth and Tom40 expression in larval muscle

Given that dMtx2 plays a conserved role in controlling the biogenesis of β-barrel proteins, we wondered whether the loss of β-barrel proteins accounts for the *dMtx2* mutant phenotypes in *Drosophila* development. The downregulation of Porin in *dMtx2* mutants does not appear to substantially contribute to the observed muscle developmental defects and pupal lethality, given that homozygous flies carrying a *Porin* deletion allele remain viable (38). In contrast, *Tom40* LOF has been shown to cause animal lethality (39–41). We expressed transgenic RNAi in muscles and found that knockdown of Tom40 or Tom22 resulted in larval lethality (Fig. 7A). When we bypassed the larval stages and activated RNAi expression in the pupal stages using *Tub-GAL80^ts^*, knockdown of Tom40 or Tom22 also caused partial or complete pupal lethality (Fig. 7B). A more careful examination of larval growth showed that knockdown of both genes in muscles caused larval growth arrest at early third instar larval stage (Fig. 7C and 7D). These results indicate that the TOM complex is essential for muscle development, and the drastic reduction of TOM complex proteins might be a major downstream pathologic pathway in *dMtx2* mutants.

**Fig. 7.**
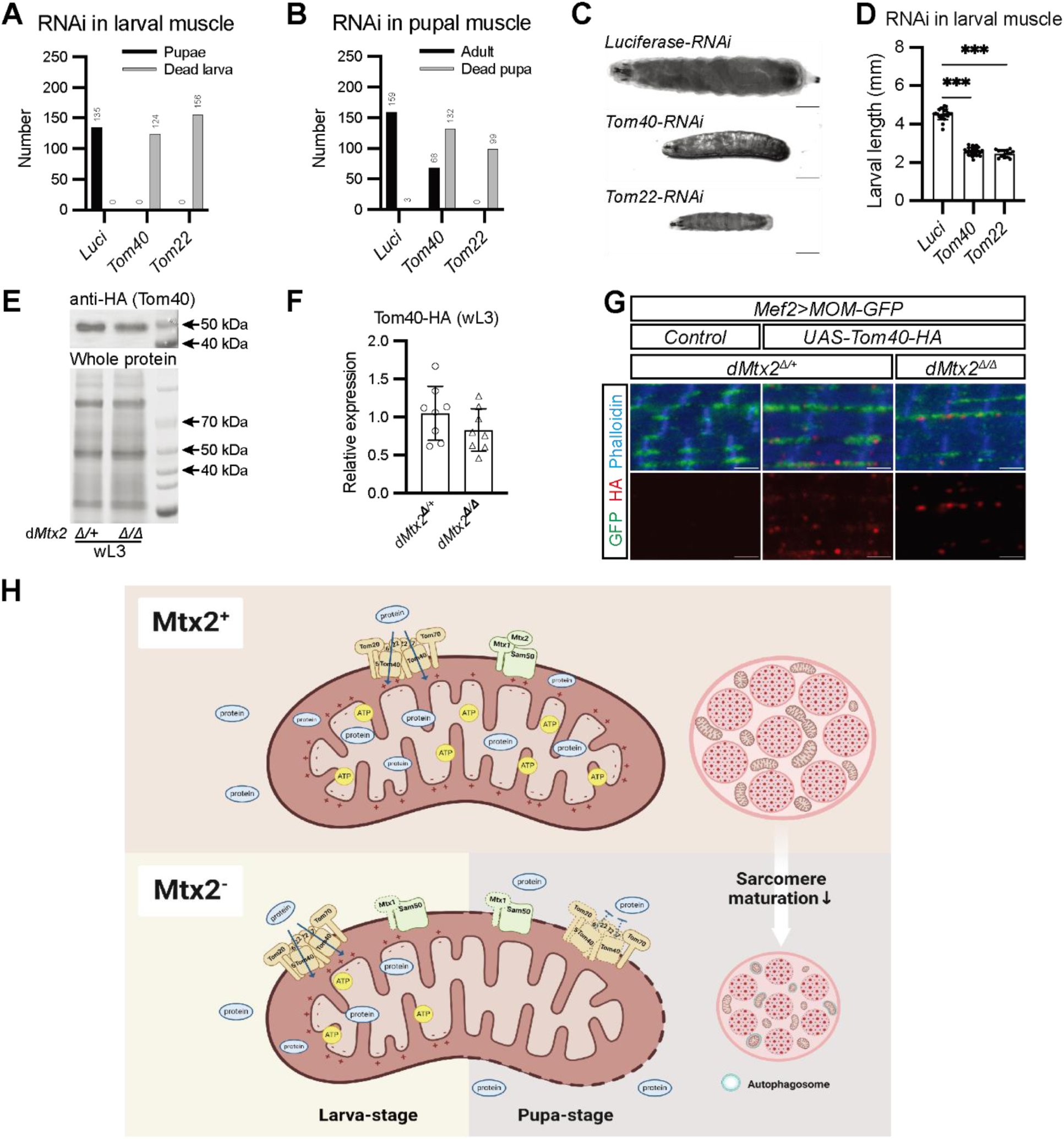
Stage-dependent roles of dMtx2 in Tom40 regulation. **(A)** Larval lethality after muscle-specific *Tom40* and *mge (Tom22)* RNAi (*Mef2-GAL4>UAS-RNAi*). **(B)** Pupal lethality after *Tom40/Tom22* knockdown specifically in pupal muscles (*Mef2-GAL4>UAS-RNAi, Tub-GAL80^ts^*, with wL3 being transferred from 18 °C to 29 °C). **(C–D)** Larval growth defects after muscle-specific *Tom40* and *Tom22* RNAi (Brown-Forsythe and Welch ANOVA test; *N*=15∼26 animals). **(E–F)** Western blot and quantification of transgenic Tom40-HA expression in wL3 muscles (Welch’s t test; *N*=8 tests). **(G)** Tom40-HA localization in larval muscles. Scale: 2 µm. **(H)** Model of dMtx2’s stage-dependent roles. The cartoon is generated on biorender.com.

However, another question arose regarding why *dMtx2* null mutants die in late pupal stages, instead of causing larval lethality. We suspected that *dMtx2* might not be required for Tom40 expression in larval muscles. Therefore, we expressed *UAS-Tom40-HA* in muscles and examined its expression in third instar larvae by Western blot analyses (Fig. 7E and 7F). Strikingly, Tom40-HA was only mildly reduced in *dMtx2* mutant larvae compared to the controls. Immunostaining results show that Tom40-HA formed puncta in larval muscles and similar Tom40 expression patterns were observed in *dMtx2* mutant animals (Fig. 7G). These results suggest that Mtx2 modulates Tom40 biogenesis differently in larval and pupal muscles. Tom40 can integrate into the MOM without dMtx2 in larval muscles, while Tom40 biogenesis is largely blocked in the absence of dMtx2 in pupal muscles.

In summary, we have revealed a stage-dependent function of dMtx2 in regulating mitochondrial β-barrel protein biogenesis and muscle development (Fig. 7H). In pupal muscles, loss of dMtx2 abolishes Tom40 integration and TOM complex assembly, leading to multifaceted mitochondrial deficits, and eventually causing muscle development arrest and animal death.

## Discussion

Autosomal recessive mutations in genes encoding components of the SAM and TOM complexes have been identified in patients with progeroid disorders (22–24, 42, 43), yet the pathological alternations leading to specific symptoms in these diseases remain largely unknown. The SAM and TOM complexes are integral to mitochondrial protein import pathways, which are essential for maintaining normal mitochondrial function in eukaryotic cells. While the protein components of these complexes have been identified and their functions extensively studied in fungi, their roles in animals are not fully understood. Specifically, how SAM and TOM complex dysfunction affects different tissues during development remain largely unexplored. Animal models are thus invaluable for elucidating the molecular and developmental mechanisms underlying these mitochondria-related progeroid diseases.

Our previous work demonstrated that *Drosophila* Mtx2 conserves its function in regulating mitochondrial transportation and morphology in neurons (30). Here, we extend our genetic investigations of *dMtx2* and uncover multiple phenotypic aspects of *dMtx2* null mutants, providing substantial evidence that dMtx2 shares conserved functions with human Mtx2. First, the expression of transgenic dMtx2 or hMtx2 successfully rescued the pupal lethality of *dMtx2* null mutants, indicating that hMtx2 can partially substitute for dMtx2. Furthermore, transgenic hMtx2 carrying MADaM patient mutations failed to rescue the lethality of *dMtx2* null mutants, confirming that these mutations are null or strong hypomorphs. Second, dMtx2 mutants display mitochondrial morphological and physiologic defects in muscle cells, including decreased mitochondrial sizes, reduced mitochondrial membrane potential and ATP production, and elevated mitophagy phenotypes similar to those observed in MADaM patient fibroblasts. Third, Mtx2 deficiency is known to cause concurrent loss of Mtx1, Tom40, and Porin in HeLa cells and MADaM patient fibroblasts (15, 22), and these proteins are also significantly reduced in *dMtx2* mutants. These findings demonstrate the similarity between dMtx2 and hMtx2 at both molecular and phenotypic levels, supporting the use of fly *Mtx2* mutations as a genetic model for studying the pathologic mechanisms of Mtx2-related diseases.

MADaM patients exhibit growth retardation, with reported cases of hypotonia (24) and cardiomyopathy (23). However, the exact mechanisms by which muscular deficits contribute to these clinical symptoms remains unknown. Tissue-specific analyses reveal that the pupal lethality observed in *dMtx2* mutants is largely attributable to muscle growth defects. Pupal muscles are formed through a series of ordered developmental events, from early myoblast proliferation to late myofibrillogenesis (44). For flight muscles, a type of striated muscles that share developmental similarities with vertebrate skeletal muscles, myofibril assembly is completed and myobifrillogenesis begins during the second day of pupal stage (45). A brief activation of dMtx2 expression in muscles on this day can support pupae eclosion. The absence of dMtx2 during this period causes permanent damages, halting sarcomere growth and maturation. The existence of this critical developmental window suggests that dMtx2 regulates muscle growth with high developmental stage-specificity. The cell-autonomous role of dMtx2 in muscle underscore the need for future clinical investigations into muscular defects and mechanistic studies in MADaM patients.

How does Mtx2 deficiency result in muscle developmental defects? Here, loss of Tom40 and impaired protein import appear to be key molecular events directly downstream of dMtx2 deficiency. The expression of Tom40 and other TOM components are significantly affected in *dMtx2* mutant pupae. Expression of Tom40 and Tom22 in muscles are essential for animal growth and survival in both larval and pupal stages, supporting the idea that disrupted TOM complex function in *dMtx2* mutants leads to subsequent mitochondrial dysfunction and muscle developmental defects. Interestingly, dMtx2 is not equally required for Tom40 expression at different developmental stages. The dMtx2 is dispensable for Tom40 expression and animal growth in larval muscles, but it becomes critical for Tom40 expression, muscle growth and animal survival in later pupal stages.

What mechanisms underlie the stage or tissue specific regulation of Tom40 expression by dMtx2? Previous studies in yeast may provide some insights. First, SAM complex components are unequally required for β-barrel protein (Tom40, VDAC and Sam50) import and folding in the MOM. In yeast, unlike Sam35 and Sam50, which are essential for cell survival, Sam37-deficient cells are viable at lower temperatures and exhibit strong growth deficits at 30 °C or higher (10, 46). Concurrently, Tom40 import and TOM complex formation are only minimally affected at low permissive temperature but are strongly impaired at non-permissive temperatures. Second, within the TOM complex, Tom6 and Tom22 play important roles in stabilizing the complex (47, 48). Overexpression of Tom6 and Tom22 can partially compensate for the loss of Sam37 in yeast under certain conditions (49, 50). Lastly, recent structural analyses reveal the architecture and folding processes of yeast SAM complexes and β-barrel proteins (8, 51, 52). Combined functional and structural results suggest that Sam37 may not directly participate in Tom40 folding, but rather modulates Sam50 activity and maintains Tom40 stability within the SAM complex. A cell’s internal state, such as its metabolic activity, may influence the importance of Sam37 to the cell’s viability. At lower temperature, yeast cells grow slowly and requires less energy supply, whereas elevated temperatures promote rapid proliferation, necessitating increased metabolic resources. Mitochondrial biogenesis and protein import must be up-regulated to meet this demand, thereby introducing potential perturbations to the assembly and stability of TOM complex. Under these stringent conditions, the modulatory function of Sam37 emerges as a critical mechanism in maintaining TOM integrity. In animals, Mtx2 may have similar functions as Sam37, so that the internal metabolic states and demands within different cells/tissues may have a role in determining their vulnerability to Mtx2 deficiency and in the generation of tissue-specific phenotypes in *Drosophila* or in MADaM patients.

Beyond the Tom40 subunit and the TOM complex, a critical question remains: what downstream molecular events underlie the observed muscle defects? Reduction of nuclear-encoded mitochondrial proteins, including mitochondrial ribosomal components and oxidative phosphorylation proteins, are detected in *dMtx2* mutants. It is well-established that mutations in mitochondrial proteins can lead to myopathy and muscle atrophy via mitochondrial dysfunctions, including reduced ATP production, disrupted mitochondrial metabolism, impaired mitochondrial calcium homeostasis, and elevated reactive oxygen species (ROS) levels (27). Such dysfunction may also trigger mitophagy and apoptosis. In *dMtx2* null mutants, a number of mitochondrial abnormalities are observed. However, our attempts to rescue mutant lethality through targeted genetic manipulations in muscles‒including expression of caspase inhibitor P35, antioxidants SOD1/SOD2, and RNAi-mediated knockdown of *Atg7* and *Atg8*‒were unsuccessful (data not shown). Intriguingly, proteomic analysis revealed substantial downregulation of both mitochondrial and cytosolic ribosomal proteins in *dMtx2* mutants (Fig. 6C), suggesting a global impairment of protein synthesis machinery. This raises the possibility that developmental phenotypes in these mutants arise not only from mitochondrial dysfunction but also from disrupted translation regulation, which may be linked reduced filament protein expression. Future studies should explore the mechanistic links between Mtx2 deficiency, protein translation alternation, and muscle-specific developmental failures.

In addition to Mtx2, Tom7 mutations have been recently identified in patients with a progeria disorder that exhibits similar manifestations as MADaM. The mutated amino acids in Tom7 are located within the Tom7-Tom40 interface, impairing interactions between these two TOM components (53, 54). Therefore, diseases associated with Mtx2 and Tom7 may converge on a common pathological mechanism involving TOM complex activity. Genetic or pharmaceutical interventions to enhance Tom40 stability and TOM complex activity may emerge as a common treatment strategy for these progeroid diseases. The tissue-specific regulation of Mtx2 highlights the complexity of mitochondrial protein import *in vivo* and underscores the importance of using model organisms to study human diseases. Our validation of *dMtx2* supports its utility as a functional model of MADaM, which can be used to elucidate the molecular and developmental mechanisms of mitochondria-related progeroid disorders and to develop potential treatment strategies.

## Methods and Materials

### Fly genetics

The following strains and genetic tools were acquired from Drosophila stock centers: *Act5C-GAL4* (BDSC#3953), *Mef2-GAL4* (BDSC#27390), *Mhc-GAL4* (BDSC#55132), *R57C10-GAL4* (BDSC#39171) (55, 56), *repo-GAL4* (BDSC#7415), 20x*UAS-FLP.D5* (BDSC#55805), *Tub-GAL80ts* (BDSC#7108), *Unc-89-GFP* (*Obscurin-GFP*) (VDRC#318326), *UAS-mito-DsRed* (BDSC#93056) (35), *UAS-mito-GFP* (BDSC#42737), *UAS-mt-Keima* (KDRC#10327), *UAS-Luciferase-dsRNA(JF01355)* (BDSC#31603), *UAS-Tom40-dsRNA(JF02030)* (Tsinghua Fly center, #2146) (41), *UAS-mge(Tom22)-dsRNA(KK108018)* (VDRC#109502) (40), *Tom40-sfGFP* (VDRC#318357), *mge-sfGFP* (VDRC#318174). *UAS-Tom40-Flag-HA* was kindly gifted by Leonie Quinn (Australian National University) (57). *UAS-MOM-GFP* was kindly gifted by Frank Schnorrer (Aix Marseille University) (34). *Tubulin-GAL4* was kindly gifted by Feng He (Zhejiang University). *Cg-GAL4* and *UAS-Tom70-HA* were kindly gifted by Chao Tong (Zhejiang University). The following mutant and transgenic lines were constructed in our lab: *dMtx2^1^*, *dMtx2^ΔDsRed^*, *dMtx2^EGFP^*, *UAS-dMtx2-EGFP*, *UAS-hMtx2-EGFP*, *UAS-hMtx2-EGFPc.294-295del*, *UAS-hMtx2-EGFPc.603delG*, and *UAS-hMtx2-EGFPc.501instT*.

### Generation of UAS transgenes of wildtype and pathogenic hMtx2

The *UAS-hMtx2-EGFP* line was created by subcloning the coding sequence (CDS) of human *Mtx2* and EGFP into pUAST-attB vector. The *dMtx2* CDS was amplified from human cDNA by PCR with primers: ATGTCTCTAGTGGCGGAAG and TGACAGCCTGCCTTTACC. *In vitro* mutagenesis was conducted by PCR with primers that embedded *hMtx2* mutation sites and by subsequent NEBuilder HiFi DNA assembly reactions (NEB).

Subcloned pUAST-attB plasmids were injected into a *Drosophila* strain carrying the PBac{yellow[+]-attP-3B}VK00002 docking site. Successful transformant was selected by the mini-white marker located within pUAST-attB vector. All fly embryo injections were carried out by UniHuaii Technology (Zhuhai, China).

### Generation of *CG9393^Myc^* knock-in fruit flies

Myc tag sequence was inserted to the 3’ end of *CG9393* CDS by CRISPR-Cas9 mediated homology recombination. Two gRNA sequences (TGCAGAGATGCGATTAGTCT and CTACGATGGCATAGATTACG) were subcloned into pCFD5 vector. A donor construct was constructed by subcloning a synthesized DNA sequence into pBluescript KS+ vector, which contained Myc coding sequence and two homolog arms (150 bp sequences adjacent to CG9393 stop codon). A 3xP3:DsRed minigene was inserted downstream to the Myc sequence in the donor construct. The gRNA and donor plasmids were transformed into *nos-Cas9* embryos and positive recombinants were selected in F1 progenies using the DsRed marker. The insertion in *CG9393* were validated by genomic DNA PCR using targeted primers (GATAGTTCTGGTTCAATGTCTTC and TATCTACGACGAGGGGATCA). The expression of Myc-tagged CG9393 was detected by Western Blot.

### Measurement of animal survival during development

Newly hatched first instar larvae were collected and reared on apple juice agar plates supplemented with yeast paste. Each plate contained 50 larvae, with 5–6 biological replicates per genotype. Larvae were transferred to fresh food plates daily until the fourth day after egg hatching, and the number of surviving larvae was recorded. Survival rate was calculated as the ratio of remaining larvae on the fourth day to the initial population. To study pupal survival rate, 50 late third-instar larvae per genotype were collected and transferred to vials with standard fly food, with 5–6 biological replicates established for each genotype. Larvae that successfully pupated were counted, and the pupariation rate was calculated as the ratio of pupae formed to the initial larval population. Pupae were maintained in standard food vials until adult eclosion. Subsequently, adult survival rate was determined by dividing the number of emerged adults by the total pupae.

For the temperature-shift regimen, parental flies were maintained at 18°C or 29°C and transferred to fresh food vials every 2–3 days. Under specific experimental protocols, flies at designated developmental stages were collected and transferred to 29°C or 18°C to modulate GAL4 activity. The initial population and the number of adults that successfully eclosed were recorded. Survival rate was calculated as the ratio of eclosed adults to the initial population.

### Measurement of *Drosophila* adult longevity

Newly eclosed female or male flies were separately collected and raised on regular food for 20 individuals per vial, with more than 4 vials for each genotype and gender. Flies were transferred to fresh food every 2–3 days. The number of flies in each vial was documented every 2 days until all flies had perished. Longevity curves were drawn based on the survival rates (survival ship) for each vial at different ages. The mean lifespan was calculated by pooling the data for all flies of each genotype and gender.

### Bright field microscopy and measurement of larval length

Larvae were fixed in 4% para-FA overnight and mounted on glass slides before they were imaged using a 5x objective lens and the DIC module under an upright microscope (Zeiss, Axio Scope A1). The collected images were analyzed using the ImageJ for measuring the larval body length.

For taking the pupal and adult photos, puparium-removed pupae and cold-anesthetized adult flies were transferred to glass slides and immediately imaged under a stereo microscope (Olympus, SZX16).

### Immunohistochemistry and fluorescent microscopy

Tissues were dissected in cold PBS and immediately fixed in PBS with 4% paraformaldehyde (PFA) and 0.3% Triton X-100 for 20 min. Fixed tissues were washed in PBS with 0.3% Triton X-100 (0.3% PBST) twice for 15 min each, and blocked in PBS with 5% normal donkey serum (NDS) and 0.3% Triton X-100 for at least 60 min. Samples were incubated for 2 days at 4 °C with primary antibodies that were prepared in blocking buffer. Samples were washed 3 times for 20 min each in 0.3% PBST before they were incubated with second antibodies for 2 days at 4 °C. After washing, tissues were stored in Fluoromount-G with DAPI, and then mounted on glass slides. All steps were performed at room temperature, unless otherwise stated. Fluorescent images were acquired with the laser-scanning confocal microscope (Leica, TCS SP8X) or epifluorescence microscope (Zeiss, Axio Scope A1). Images were processed using ImageJ (GraphPad). Please refer to Table S1 for the antibodies used in this study.

### Electric microscopy

Pupae were collected 96 hours after puparium formation. The pupal case was removed, and the abdomen and head were manually dissected using a microtome blade. Thoraces were fixed overnight at 4°C in 0.1 M phosphate buffer (PBS, pH 7.0) containing 4% paraformaldehyde and 2.5% glutaraldehyde. After fixation, thoraces were post-fixed with 1% osmium tetroxide (OsO₄) in 0.1 M PBS for 1 hour at room temperature. Samples were then washed three times with 0.1 M PBS (5 minutes per wash), followed by incubation in 1% uranyl acetate (in H_2_O) at 4°C for 2 hours. Dehydration was performed by a series of 10 min incubations in 50%, 70%, 80%, 90%, 100% ethanol and finally by propylene oxide thrice. The dehydrated thoraces were embedded in Epon resin and polymerized at 60°C for 48 hours. Ultra-thin sections (80 nm) of the sample blocks were cut by a Leica UC 7 microtome (Leica, Vienna, Austria) with diamond knife (Diatome, Switzerland). Imaging was conducted on an HT7700 transmission electron microscopy (Hitachi, Japan) operated at 80 kV, and micrographs were captured with a Gatan 830 CCD camera.

### Live imaging of pupal DLMs

Pupae were dissected and obtained in HL3 buffer (70 mM NaCl, 5 mM KCl, 20 mM MgCl_2_•6H_2_O, 10 mM NaHCO_3_, 1.5 mM CaCl_2_, 115 mM Sucrose, 5 mM Trehalose, 5 mM HEPES) for live imaging using an inverted confocal microscope.

For analysis of ΔΨm, dissected semi-thorax were plated in 96-well plates (CellCarrier, PerkinElmer) and subsequently incubated in 40 nM TMRM (Beyotime, C2001S-1) with or without the presence of 10 μM CCCP (Beyotime, C2001S-3) for 30 min at room temperature. After incubation, tissues were transferred into 5 nM TMRM buffer on a glass slide during image acquisition. TMRM fluorescence (Ex: 550 nm/Em: 570–620 nm) images were acquired using oil objective (60x, NA 1.42) of confocal (Olympus, FV3000). TMRM fluorescence intensity was then measured and averaged using ImageJ software. At least two DLMs per animal and more than five animals for each genotype were imaged. The average value from each animal was used as one datapoint for statistical analysis performed by Prism 9 (GraphPad).

For mt-Keima analysis, confocal Z-stacks of the pupal DLM were acquired using Leica SP8 system with a 63x oil-immersion lens (NA 1.40). The sample was excited using 488 or 552 nm lasers and the emission light (600-700 nm) was separately collected. The Z-stacks (24.05 μm x 24.05 μm x 10 μm) were projected to a single image along Z-axis using the ImageJ software. Mitochondria of sizes > 0.04 μm^2^ in the 552 nm channel were counted. At least two DLMs per animal and more than five animals for each genotype were imaged. The average value from each animal was used as one datapoint for statistical analysis performed by Prism 9 (GraphPad).

For live imaging of MOM-GFP and mito-DsRed, MOM-GFP (Ex: 488 nm/Em: 500-541 nm), and mito-DsRed fluorescence (Ex: 555 nm/Em: 570–620 nm) images were acquired sequentially, by Leica SP8 using an oil objective (63x, NA 1.40). The analysis was performed by ImageJ/threshold to detect mitochondria positive region in the MOM-GFP channel. The fluorescence intensity of the same region in mito-DsRed channel was then measured.

### SDS-PAGE and Western blot

SDS–PAGE and Western blot were conducted following standard protocols. Basically, 5 pupae or larva were homogenized in 200 μL of RIPA lysis buffer (Beyotime Biotechnology, P0013B), 2 μL of PMSF, and 2 μL of protease inhibitor cocktail (MCE, HY-K0010). The lysate was centrifugated at 4°C at 13500 rpm for 10 minutes and 80 μL of the supernatant was mixed with 20 μL of 5 × Loading buffer (Beyotime, P0015). Samples were loaded onto a 10% TGX stain-free SDS–PAGE (Bio-Rad, 1610183), and the quantities of total proteins were analyzed by Bio-Rad ChemiDoc MP imaging system after electrophoresis. For Western blot, Proteins were transferred onto a PVDF membrane. The blot was blocked with 5% BSA and was incubated with primer antibodies prepared in blocking buffer overnight. After washing, the blot was incubated with HRP-conjugated secondary antibodies and imaged by using Bio-Rad ChemiDoc MP imaging system after ECL reaction. Please refer to Table S1 for the antibodies used in this study.

### Quantitative proteomics analysis of *Drosophila* pupae

#### Protein extraction

Frozen pupae (about 100 mg) were transferred into low protein binding tubes and lysed with 300 µL lysis buffer supplemented with 1mM PMSF. Then samples were further lysed with sonication. The parameters were set as 1s /1s intervals and 80 W for 2 min. After sonication, the samples were centrifuged at 12000 rpm for 10 min at 4°C to remove insoluble particles, repeat once to further exclude precipitation. Protein concentration was determined by BCA assay. The protein samples were aliquoted to store at-80°C.

#### Protein digestion

According to the measured protein concentration, take 50 μg protein from each sample, and dilute different groups of samples to the same concentration and volume. Add 25mm DTT of the corresponding volume into the above protein solution to make the DTT final concentration about 5 mM, and incubate at 55°C for 30 min. Then add the corresponding volume of iodoacetamide so that the final concentration was about 10 mM, and place it in the dark for 15 min at room temperature. Then 6 times of the volume of precooled acetone in the above system to precipitate the protein, and place it at −20 °C for more than four hours or overnight. After precipitation, take out the sample and centrifuge at 8000 g for 10 min at 4 °C for collecting the precipitate. According to the amount of protein, add the corresponding volume of enzymolysis diluent (protein: enzyme = 50:1 (m/m), 100 μg of protein add 2 μg of enzyme) to redissolve the protein precipitate, then the solutions were incubated for digestion at 37°C for 12 h. Finally, samples were lyophilized or evaporated after enzymolysis.

#### Label

For TMT labelling, the lyophilized samples were resuspended in 30 μL 100 mM TEAB and Labeling reaction in a 1.5 mL Ep tube. 20 μL acetonitrile were added to TMT reagent vial at room temperature. The centrifuged reagents were dissolved for 5 min and mixed for centrifugation and repeat this step once. Then 10 μL of the TMT label reagent was added to each sample for mixing. The tubes were incubated at room temperature for 1 h. Finally, 5 µL of 5% hydroxylamine were added to each sample and incubated for 15 min to terminate reaction. The labeling peptides solutions were lyophilized and stored at −80°C.

#### Liquid chromatography-mass spectrometry

The Proteomic data analysis was performed by Shanghai Luming biological technology co., LTD (Shanghai, China). RP separation was performed on an 1100 HPLC System (Agilent) using an Agilent Zorbax Extend RP column (5 μm, 150 mm × 2.1 mm). Mobile phases A (2% acetonitrile in HPLC water) and B (90% acetonitrile in HPLC water) were used for RP gradient. The solvent gradient was set as follows: 0∼8 min, 98% A; 8∼8.01 min, 98%∼95% A; 8.01∼30 min, 95%∼80% A; 30∼43 min, 80∼65% A; 43∼53 min, 65∼55% A; 53∼53.01 min, 55∼10% A; 53.01∼63 min, 10% A; 63∼63.01 min, 10∼98% A; 63.01∼68 min, 98% A. Tryptic peptides were separated at a fluent flow rate of 300 μL/min and monitored at 210 nm. Samples were collected for 8-54 minutes, and eluent was collected in centrifugal tube 1-15 every minute in turn. Samples were recycled in this order until the end of gradient. The separated peptides were lyophilized for mass spectrometry.

All analyses were performed by a Q Exactive HF mass spectrometer (Thermo, USA) equipped with a Nanospray Flex source (Thermo, USA). Samples were loaded and separated by a C18 column (Acclaim PepMap RSLC, 50 cm × 75 µm) on an EASY-nLCTM 1200 system (Thermo, USA). The flow rate was 300 nL/min and linear gradient was 45 min (0∼4 min, 8-11% B; 4∼36 min, 11-45% B; 36∼39 min, 45-100% B; 39∼45 min, 100% B. mobile phase A = 0.1% FA in water and B = 0.1% FA in ACN). Full MS scans were acquired in the mass range of 350-1500 m/z with a mass resolution of 45000 and the AGC target value was set at 3e6. The 20 most intense peaks in MS were fragmented with higher-energy collisional dissociation (HCD) with collision energy of 32. MS/MS spectra were obtained with a resolution of 3000 with an AGC target of 2e5 and a max injection time of 40 ms. The Q Exactive HF dynamic exclusion was set for 30.0 s and run under positive mode.

#### Database search

ProteomeDiscoverer (v.2.4.1.15) was used to search all of the raw data thoroughly against the Uniprot Drosophila melanogaster database. The main parameters were set as following:

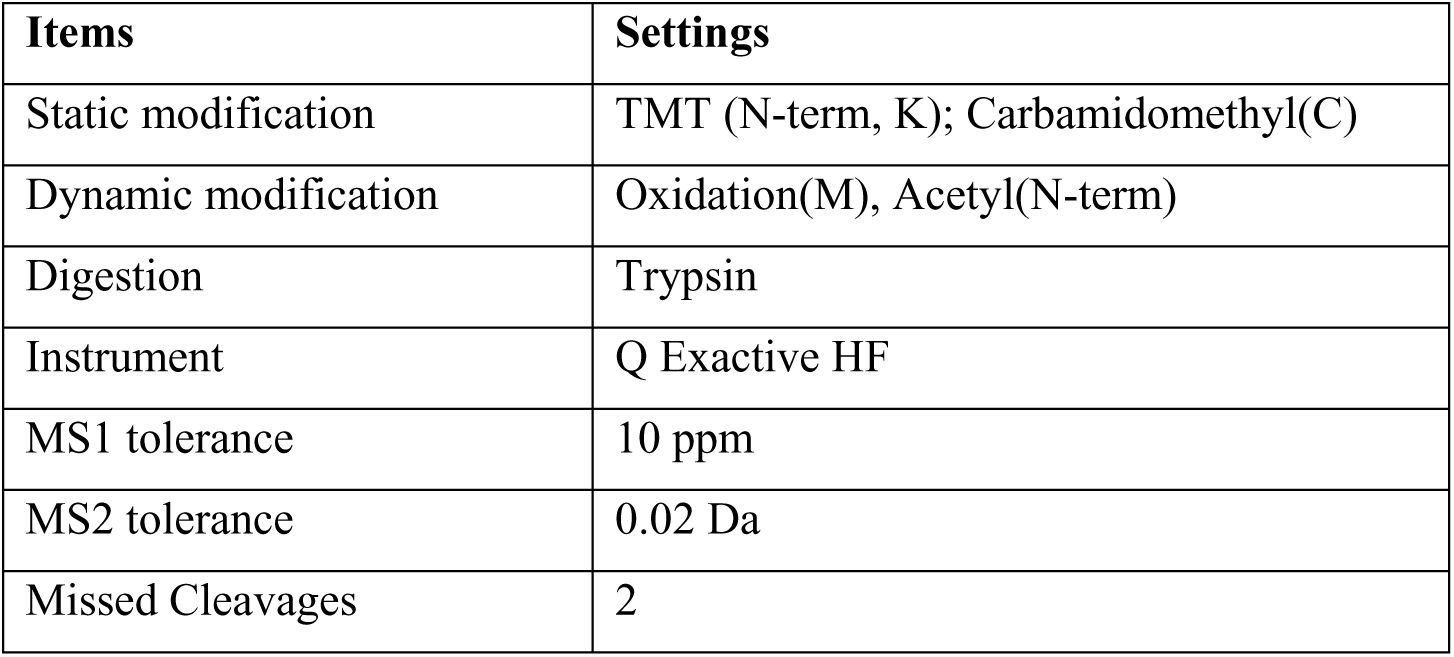

### Statistical analyses

A total of 5874 proteins expressed were identified. Differentially expressed proteins (DEPs) were identified using thresholds of |log2(fold change)| > 0.585 (equivalent to fold change (FC) ≥ 1.5 or ≤ 1/1.5) combined with a statistical significance threshold of *P* < 0.05. The log2(FC) was calculated as the mean expression value of the homozygous group minus that of the heterozygous group. Then we found 35 upregulated and 110 downregulated proteins in dMtx2 mutant homozygous compared with dMtx2 mutant heterozygous. Annotation of all identified proteins was performed using GO (http://www.blast2go.com/b2ghome; http://geneontology.org/) and KEGG pathway (http://www.genome.jp/kegg/). DEPs were further used for GO and KEGG enrichment analysis. Protein-protein interaction analysis was performed using the String (https://string-db.org/).

### Statistics

Statistical analyses were performed by using the Prism 9 (GraphPad). To assess the statistical significance of the observed differences, we employed unpaired t-tests for comparisons between 2 groups, one-way ANOVA for evaluating group differences in single-factor experiments, and two-way ANOVA for more complex interactions within 2 independent variables. The critical thresholds for significance were established as follows: \**P* < 0.05, \*\**P* < 0.01, and \*\*\**P* < 0.001. Details of the specific statistical tests employed can be found in the respective figure legends. The details of statistical analyses are summarized in Table S2. The graphical data are presented as the mean ± standard deviation (SD) to provide an overview of the central tendency and variability. *N* equals to the number of animals or tests as indicated in figure legends.

## Acknowledgements

We thank Feng He, Leonie Quinn, Frank Schnorrer, Chao Tong, Xuan Guo, the Bloomington Stock Center, the Vienna Drosophila Resource Center, the Korea Drosophila Resource Center and the Tsinghua Fly center for fly stocks. We thank Min Jiang for insightful discussions. Electron microscopy imaging was performed on the core facility of the Life Sciences Institute Zhejiang University with the help from Ziyi Kang. Confocal imaging was conducted on the Core Facilities of Children’s Hospital, Zhejiang University School of Medicine with technical support from Yanhua Bi, Yusang Dong, Fangqing Wang and Chunchun Zhi.

## Author contributions

Conceptualization: J.M and X.X.

Methodology: X.S, W.S, and X.X.

Investigation: X.S, X.F, L.L., Y.R., W.L.

Visualization: X.S, X.F., Y.R. X.X

Supervision: W.S., J.M., and X.X.

Writing: X.S., J.M. and X.X.

## Data and materials availability

Raw mass spectrometry data have been uploaded to iProX with the accession number of IPX0011604000. Protein abundance data for the proteomics are provided in Data S1. Statics analysis details are listed in Table S2. All raw images and fly strains generated in this study are available upon reasonable request.

## Supplemental figures

**Fig. S1-related to Fig. 1.**
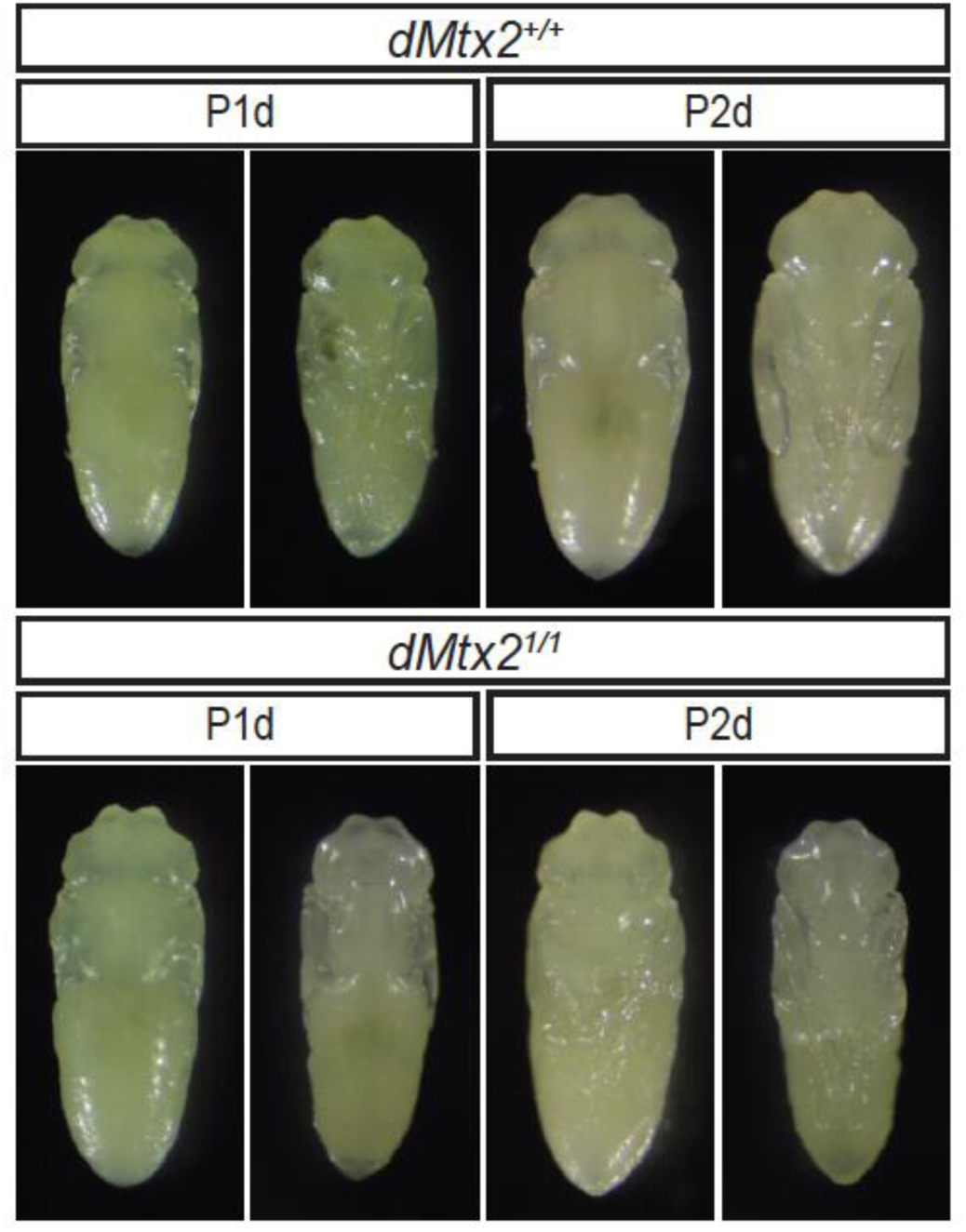
*Drosophila Mtx2* is dispensable for early pupal morphogenesis. Morphology of control and mutant pupae in early stages.

**Fig. S2-related to Fig. 1.**
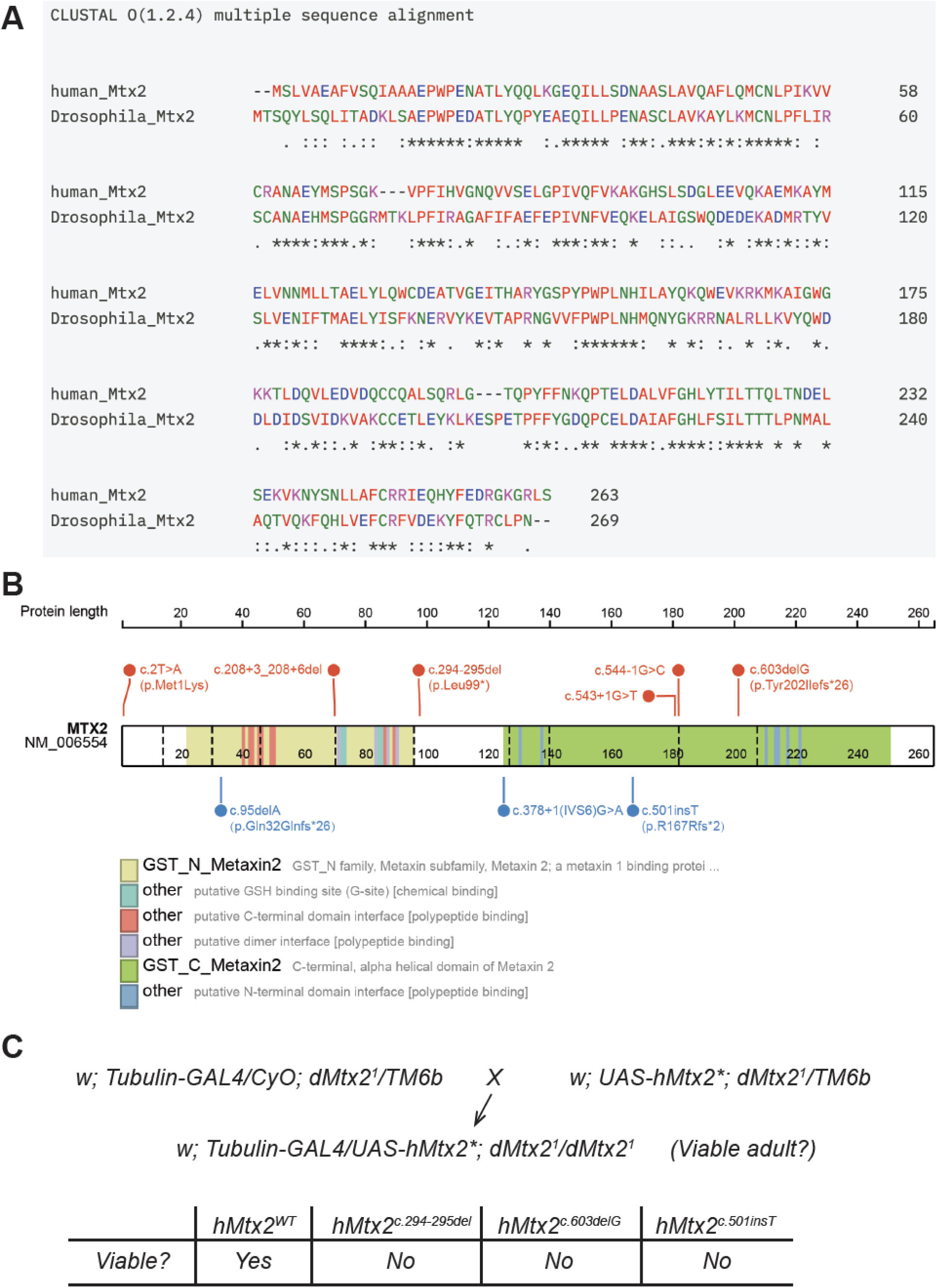
Pathogenic hMtx2 variants fail to rescue *dMtx2* mutant lethality. **(A)** Sequence alignment of human and *Drosophila* Mtx2. **(B)** Domain structure of hMtx2 with MADaM-associated mutations highlighted. **(C)** Rescue assay testing pathogenic *hMtx2* variants. The rescue effect was assessed by the emergence of mutant adults.

**Fig. S3-related to Fig. 3.**
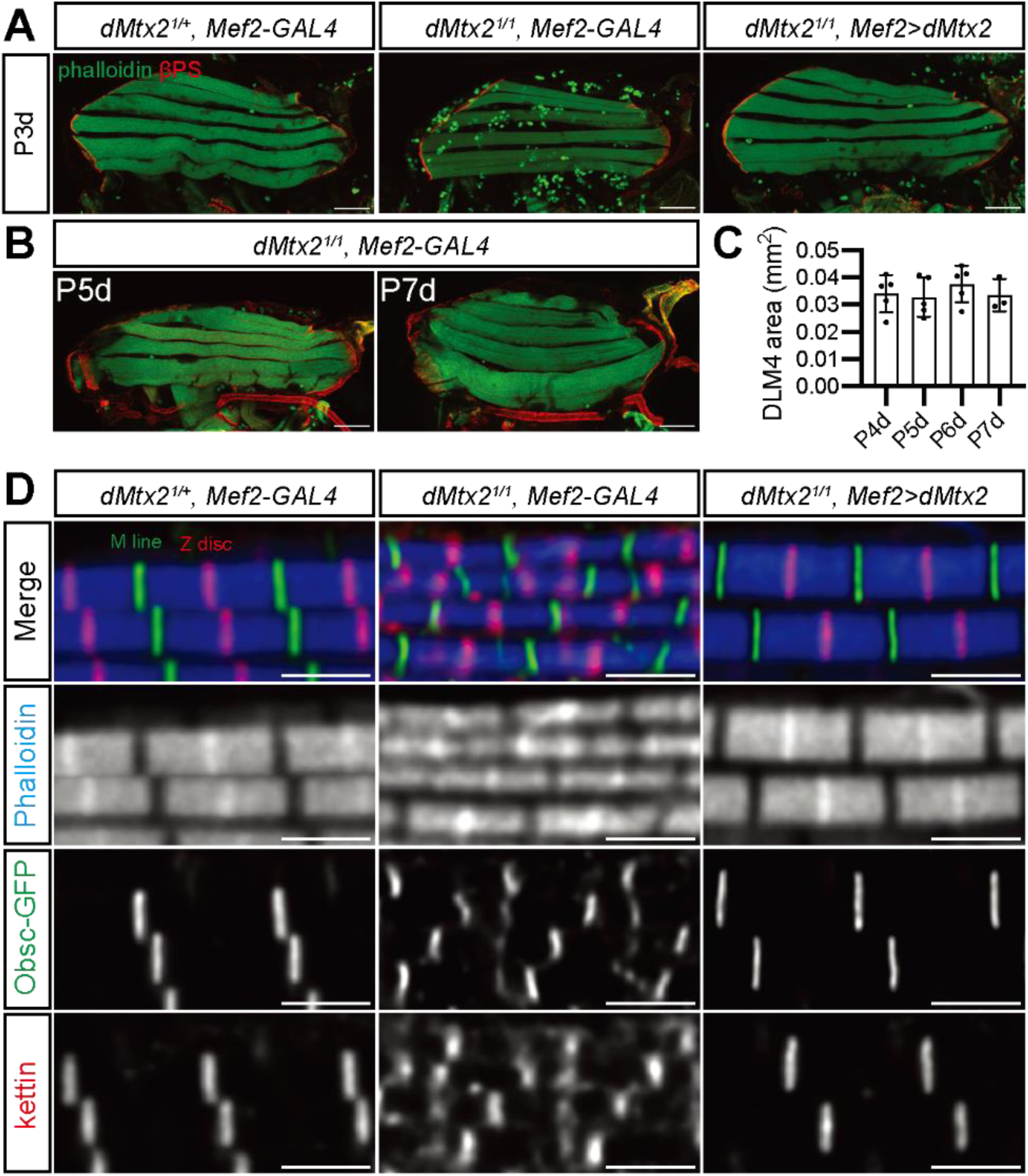
Muscle growth defects are revealed in *dMtx2* mutant pupae. **(A)** DLM staining at 3 days after pupal formation. Scale bars: 100 µm. **(B-C)** DLM staining (B) and DLM4 size measurements (C) during extended pupal stages (One-way ANOVA, Dunnett test; *N*=3∼5 animals). **(D)** Sarcomere structures labeled with phalloidin (blue), Obscurin-GFP (M-line, green) at 4 days after pupal formation, and kettin (Z-disc, red). Scale bars: 2 µm.

**Fig. S4-related to Fig. 5.**
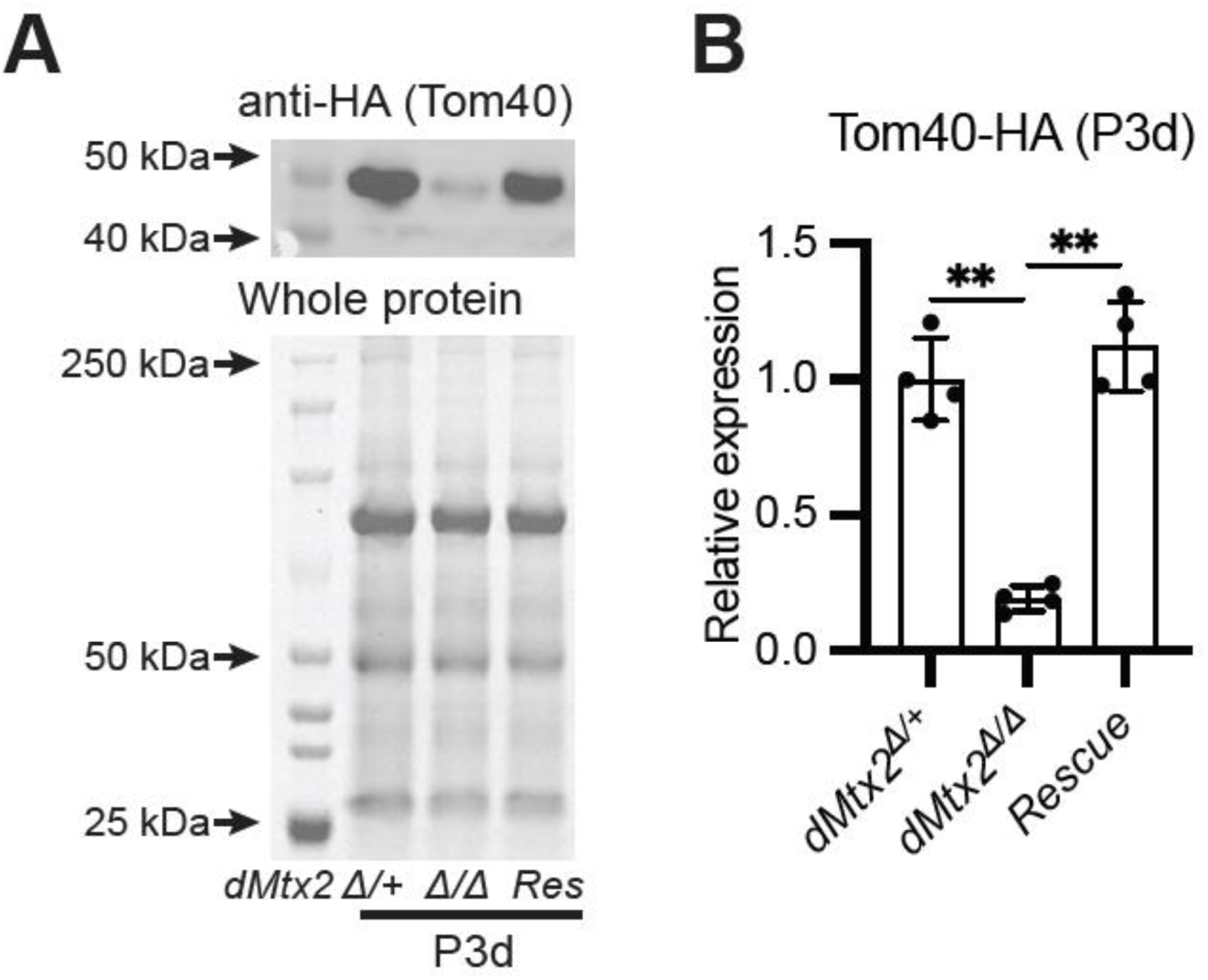
Rescue of Tom40 expression in dMtx2 mutant. Western Blot (A) and quantification (B) of transgenic Tom40-HA expression in P3d muscles (Brown-Forsythe and Welch ANOVA test; *N*=4 tests).

**Fig. S5-related to Fig. 6.**
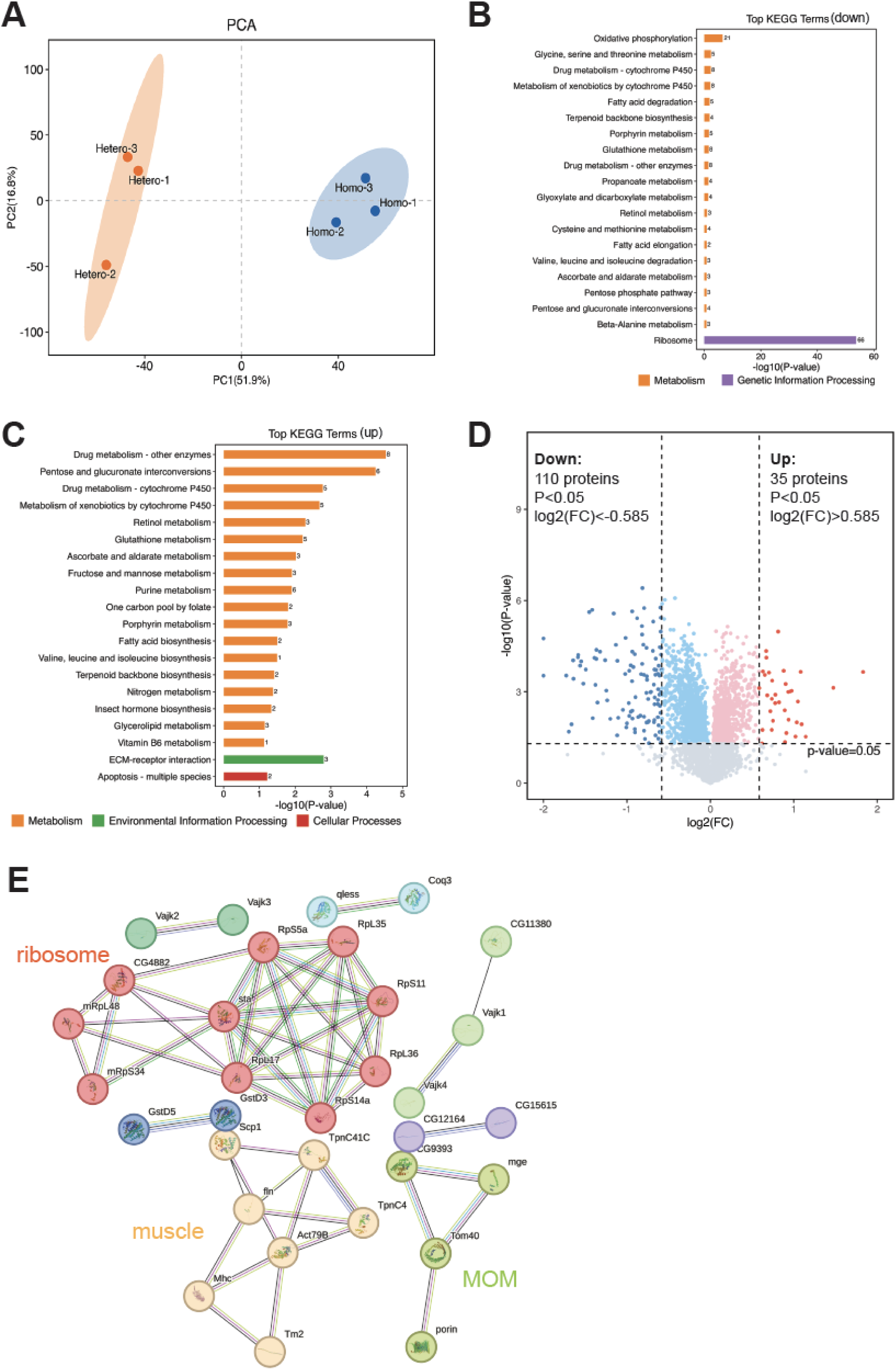
Bioinformatic analyses of the proteomic data. **(A)** Principal component analysis (PCA) of all six samples. **(B-C)** KEGG analyses of the downregulated (B) and upregulated DEPs identified by low-stringency thresholds (|fold change(FC)|>1.2, *P*<0.05). **(D)** A volcano plot of the DEPs using high-stringency thresholds (|fold change(FC)|>1.5 or |log2(FC)|>0.585, *P*<0.05). **(E)** STRING analysis showing the interaction networks of the downregulated DEPs identified in panel D.

**Fig. S6-related to Fig. 6.**
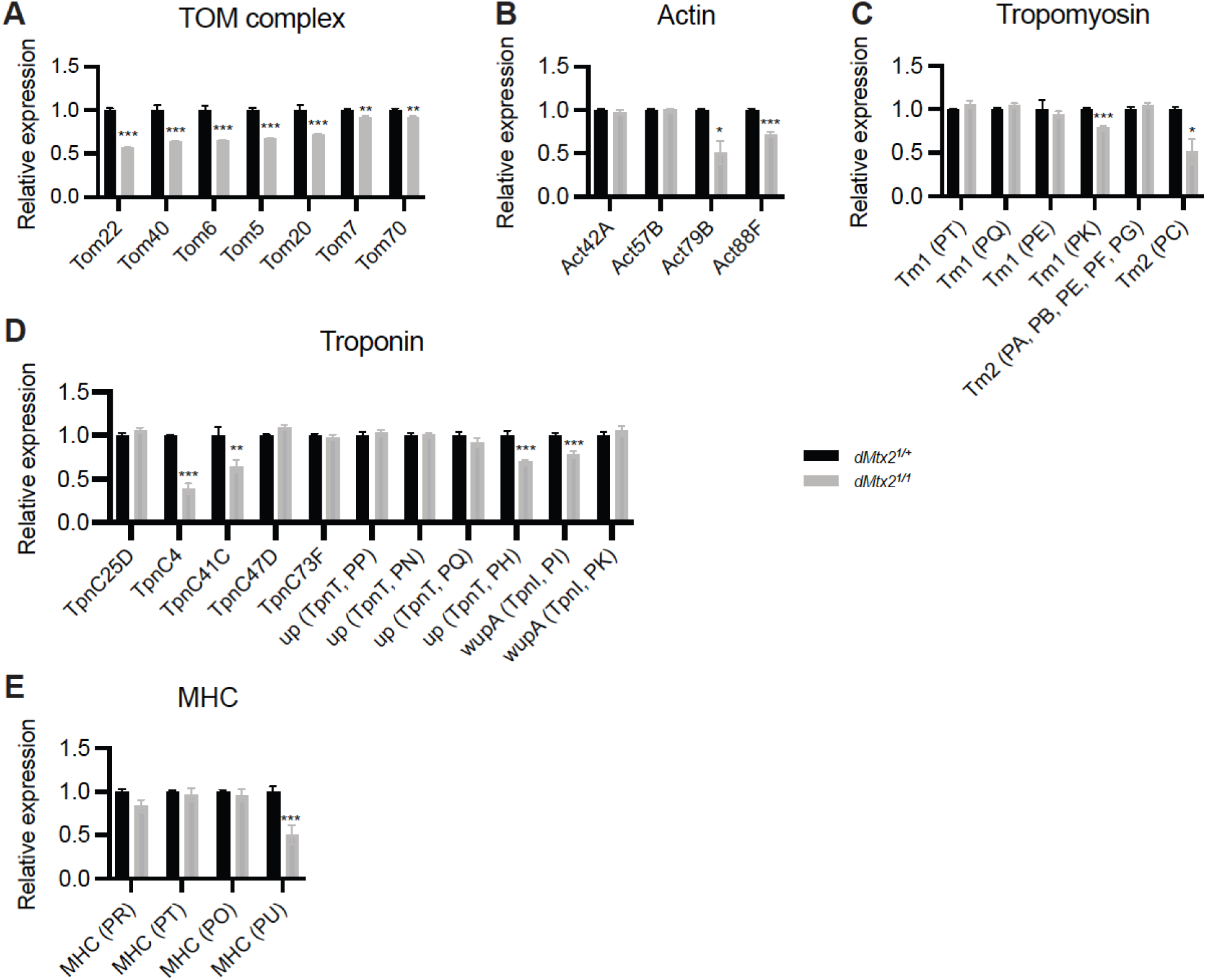
Bioinformatic analyses of the proteomic data. Bar graphs of TOM complex protein (A), actin (B), tropomyosin (C), troponin (D) and myosin heavy chain (MHC) (E) levels detected by MS/MS.

